# Epithelial Reprogramming and Transition during Pulmonary Bioengineering

**DOI:** 10.64898/2026.01.24.701406

**Authors:** Satoshi Mizoguchi, Vi Lee, Hahram Kim, Sophie E. Edelstein, Nuoya Wang, Maria Tomas Gracia, Colten Danelski, Connor Haynes, Rachel Rivero, David Stitelman, Tomohiro Obata, Allison M. Greaney, Tomoshi Tsuchiya, Themis R. Kyriakides, Naftali Kaminski, Micha Sam Brickman Raredon

## Abstract

Recent research has emphasized the critical role of cell state transitions in tissue homeostasis. In lung biology, transitional cells are recognized as a feature of tissue-scale processes during both normal physiology and disease. The precise way that transitional cell states emerge and are regulated remains to be determined. Engineered tissues, built in a laboratory through bioengineering approaches, allow detailed study of cellular states that are not commonly found in native biology, and allow opportunities to directly induce and manipulate cellular transitions. The following study explores and characterizes epithelial cell states that emerge via cellular reprogramming in a tissue engineering context.

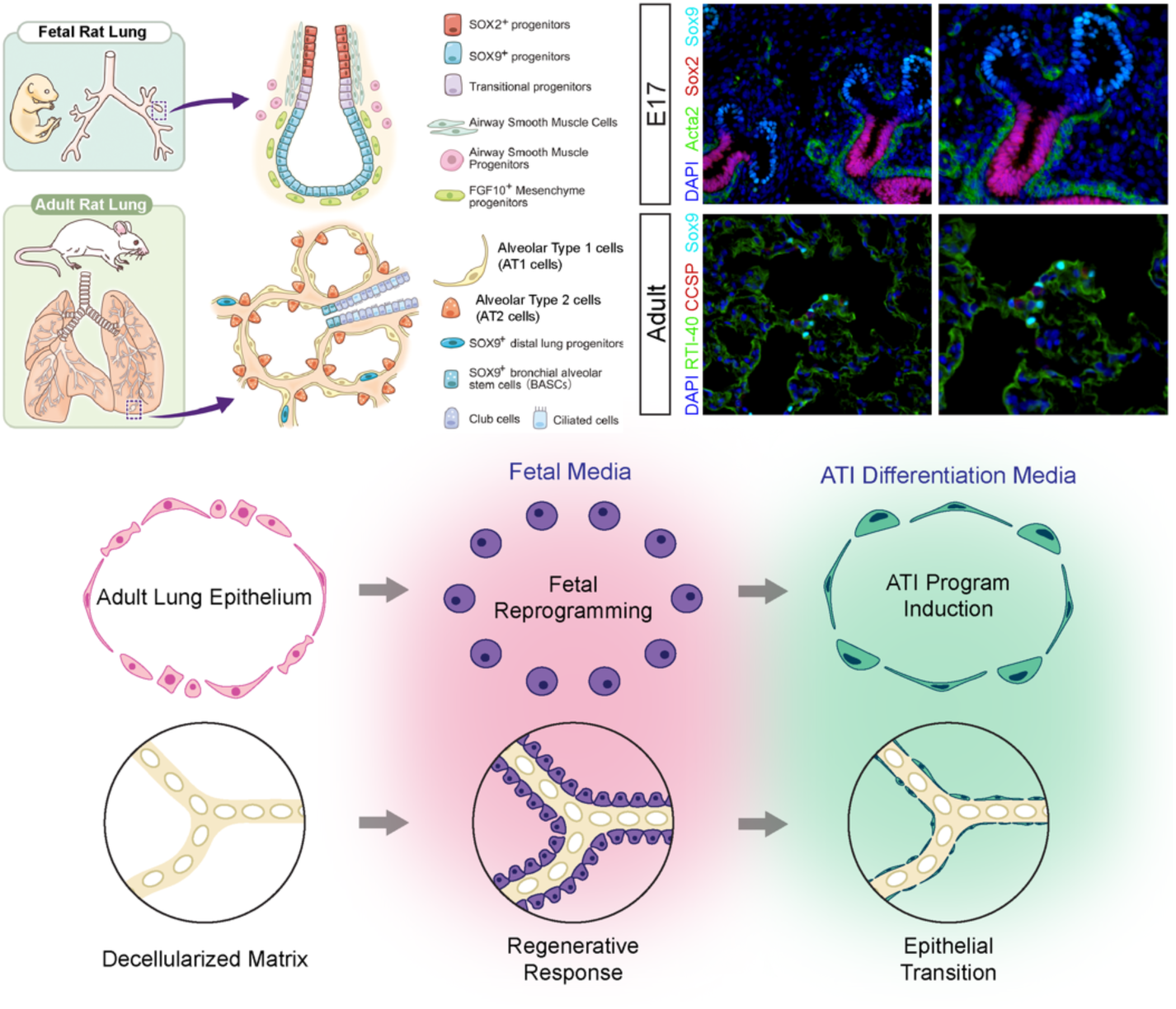

## Background

Cellular transition between cell states is fundamental to tissue biology (*1*). Cells within tissues generally reside within highly structured biochemical milieus which constrain genetic regulatory network activation and thereby canalize gene expression and resulting cellular phenotype (*2, 3*). In the lung, precisely structured cell-to-cell signaling niches maintain this cellular canalization during homeostasis and guide cellular transition during both homeostasis and repair (*4*). In disease processes, signaling niches are disrupted and cellular transition is arrested or perturbed, leading to improper tissue homeostasis and the development of pathology (*5–8*). Understanding the principles of cellular transition and mapping causal relationships between extracellular milieu and intracellular state, is of utmost importance to tissue biology and pulmonary regenerative medicine.

The study of native tissues taken from animals, both healthy and diseased, has formed the foundation of biology and medicine. However, when we study in vivo tissues, we end up only studying cell states, and sets thereof, which are found during in vivo processes. There are many living, viable, cell states which do not exist within native biology. These include cultured cells (*9–11*), cells within organoids (*12–14*), and cells within perfused and engineered tissues (*15–17*). These cell states, while often highly different from cells found in the body, can provide significant insight into the fundamental nature of cell state regulation when studied computationally (*18–20*). Bioengineering allows researchers to manipulate cell states in a deliberate fashion, and therefore, engineered tissue biology provides a powerful window into the processes and control parameters for cellular transition.

In the present study, we profile the cellular and molecular profile of a specific pulmonary epithelial reprogramming and transition trajectory that we have deliberately induced as part of ongoing bioengineering efforts. We show that by placing a heterogeneous primary pulmonary epithelial population within an extracellular milieu designed to mimic the epithelial microenvironment found in fetal lung, we induce de-differentiation, reduce cellular heterogeneity and stabilize a biphasic population of reprogrammed epithelial cells which display marked fetal character. Then, by changing the extracellular milieu to one which is designed to directly activate YAP, we induce ATI-cell associated gene expression, driving the cells directly into a transitional cell state which displays many of the transcriptional and phenotypic features known as hallmarks of transitional cells found in pulmonary fibrosis (IPF) and chronic obstructive pulmonary disease (COPD). We find that complete ATI-transition does not occur in the absence of additional alveolar niche cell types and/or on the timescale in which we have performed our experiments. Our findings confirm the ability of extracellular cues to regulate cell state and show that cells within bioengineered tissues can be driven into transitional states that highly mimic those seen in human clinical disease.

## Results

### Direct reprogramming of adult lung epithelium

We hypothesized that adult lung epithelium might be susceptible to direct developmental reprogramming if cultured in a fetal extracellular milieu. To test this hypothesis, we dissociated native adult rat lung as previously reported (*4*) and enriched the resulting single-cell suspension for epithelial cells (**Figure 1A**). These cells were cultured in a media originally designed to maintain fetal cellular states (Fetal Growth LPM-3D Media adapted from (*21*), see Methods for detailed composition)(**Figure 1B**). Within this culture media, epithelial cells proliferated generously, forming multi-layered topologies and spheroids in 2D culture (**Figure 1C**). By P3, the cells began to change character, and the morphology became more irregular and less polygonal. By P4 to P5, the cells still proliferated but no longer formed complex topologies. The cells were passaged up to P12. RT-qPCR data was collected for cells in passages P0 through P5. qRT-PCR found specific genes of interest *Trpm5*, *Sox9*, and *Pou2f3* to increase in bulk expression while *Dclk1* expression decreased (**Figure 1D-G**). Cytospin slides of P0, P3, and P5 cells were prepared (n=3, biological replicates) and stained for *Abca3*, *Ager*, *Epcam*, *Krt5*, *Sox2*. Whole-slide stitched immunofluorescence images were captured, and images were segmented and processed using simpleSeg in R (*22*) (see Methods) to yield unbiased single-cell protein expression values (**Figure 1H, Right**). Intensity thresholds were titrated for each marker to bin positive and negative cells (**Figure 1H**, **Left**). Output data were used to estimate population-scale marker positivity at each passage (**Figure 1I**). These data showed a significant decrease in *Abca3* expression (indicating loss of ATII- cell character), concurrent with an increase in *Krt5* (gain in basal-cell character) and gain in *Sox9* (gain in fetal character) from P0 up to P5.

**Figure 1.**
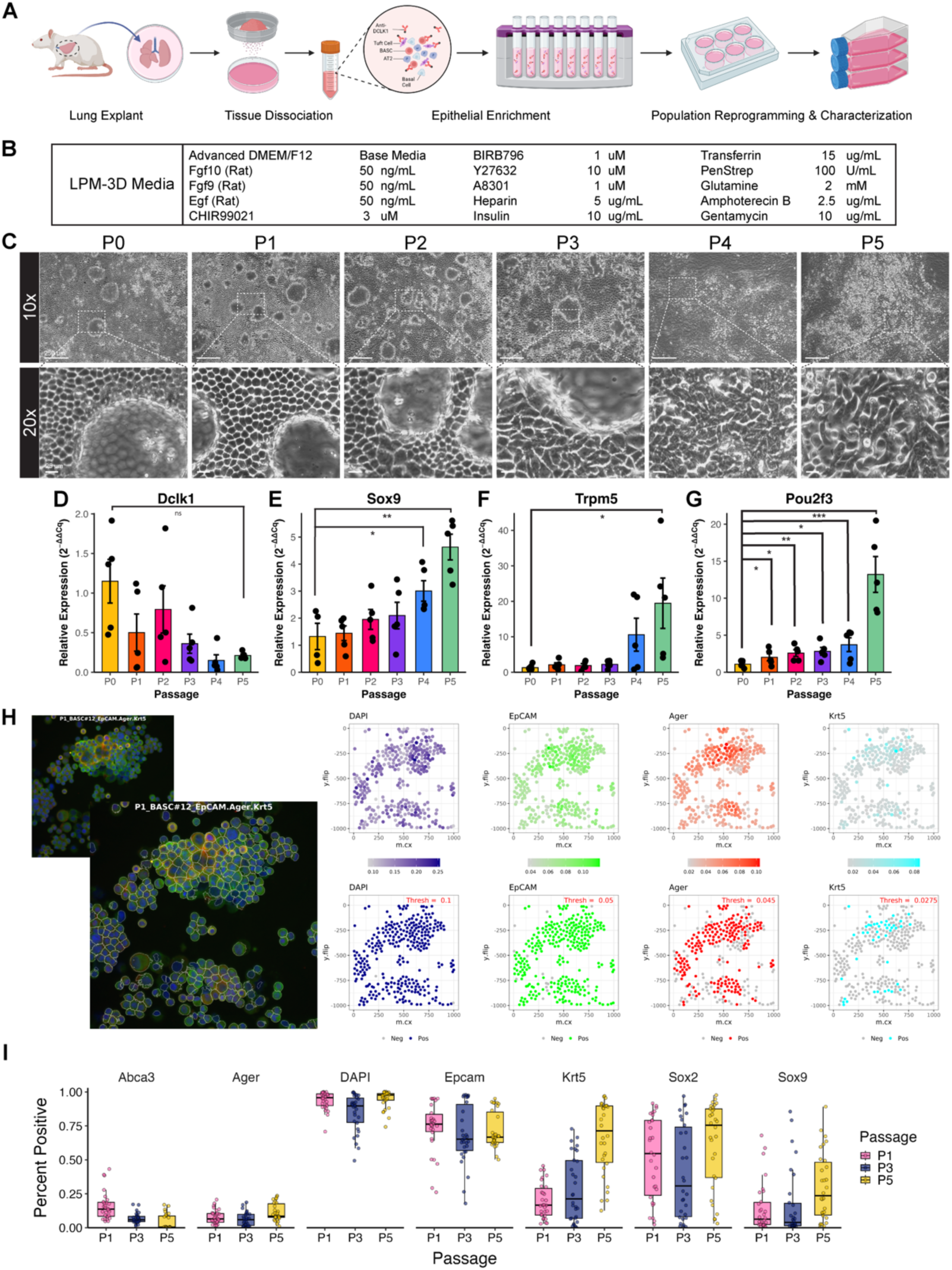
Reprogramming of adult lung epithelium. **A)** Rat lungs were explanted, the tissue dissociated, the single-cell suspension enriched for epithelial cells and profiled via scRNAseq and then passaged in fetal reprogramming medium (LPM-3D). B) LPM-3D components, which include fetal growth factors acting on epithelial cells and small molecules activating canonical WNT signaling and inhibiting both TGFβ and ROCK pathway transduction. **C)** Brightfield microscopy of cells in LPM-3D Media for Passages 0 through 5 in 2D Culture. The cells displayed a uniform, polygonal morphology and proliferate to over-confluence in earlier passages, then began to adopt a more irregular morphology with less proliferative character in later passages. Scale bars (top row, 10x): 250 µm. Scale bars (bottom row, 20x): 25 µm. **D-G)** Quantitative RT-PCR analysis of *Dclk1* (D), *Sox9* (E), *Trpm5* (F), and *Pou2f3* (G) expression across passages, normalized to β-actin and plotted as relative expression (2 ^-Δ^ ^ΔCt^). Points represent individual biological replicates, and bars represent mean ± SEM. Pairwise statistical comparisons between passages were performed using two-tailed, unpaired Student’s t-tests on ΔΔCt values, with significance indicated as *p < 0.05, **p < 0.01, and ***p < 0.001. Data reveals an increase in *Sox9* expression starting at P3 and increasing significantly by P5. **H)** Schematic of cytospin cell segmentation and single-cell protein quantification. Fluorescent microscopy images were processed using EBImage and SimpleSeq in R to quantify channel-associated relative marker expression. Data was quantified after channel intensity normalization, and thresholds were titrated for each channel in order to assign each marker in each cell as either positive or negative. **I)** Boxplots of cytospin-based single-cell marker positivity across passages (P1, P3, and P5), shown as the fraction of cells positive for each marker. Each box represents the distribution of per-sample percent-positive values, with overlaid jittered points indicating individual biological replicates. Notable trends include an increase in *Krt5* and *Sox9* expression, and a decrease in *Abca3* and *Sox2* expression. DAPI is shown as a nuclear control.

### Characterization of reprogrammed epithelium confirms fetal character

To confirm that all of the cells we were studying were indeed epithelium, and to profile their genetic expression, we turned to single-cell RNA sequencing (scRNAseq). We captured cells from our starting epithelium-enriched population before culture (n=3 biologic replicates), Passage 0 in LPM-3D (n=3 biological replicates), Passage 3 (n=3 biological replicates), and fetal lung to use as a control. We sequenced each batch of cells to a target read depth of 50,000 reads/cell and processed the resulting data through iterative cleaning, annotation, and analysis (see Methods). **Supplemental Table 1** shows data acquisition and QC metrics for all samples. We were able to identify analogous cell states in each experimental condition, which nonetheless showed great transcriptional difference at each time point. **Supplemental Figure 1** show individual condition embeddings, clustering, annotations, intra-condition cell state markers and global cross-condition transcriptomic markers for each of these 4 experimental arms. Our findings fundamentally revealed that culture in fetal media induces a loss of population-scale cellular heterogeneity and a loss of differentiated adult character (**Figure 2A-H**). Although several cell states were still recognizable as similar to canonical native cell types, none displayed fully differentiated character. The observed decrease in cellular differentiation and drop in population heterogeneity was accompanied by increased transcription of many genes which mark fetal epithelium (**Figure 2I-J**). Reprogramming culture caused transcriptional upregulation of *Wnt7b* (*23*), *Tgfb2* (*24*), *Efnb2* (*25*), and *Lamc1* (*26*), all of which are highly expressed by developing epithelium (**Figure 2I-J**). These changes occur in tandem with genes regulating cellular proliferation and inhibiting apoptosis (*Mki67*, *Birc5*, *Topbp1*) which also show high transcription during development (*27–30*)(**Figure 2J**). These findings demonstrated to us that we had reprogrammed adult epithelial cells to a fetal-like state, inducing many, though certainly not all, genetic programs seen within epithelial cells during normal lung development.

**Figure 2.**
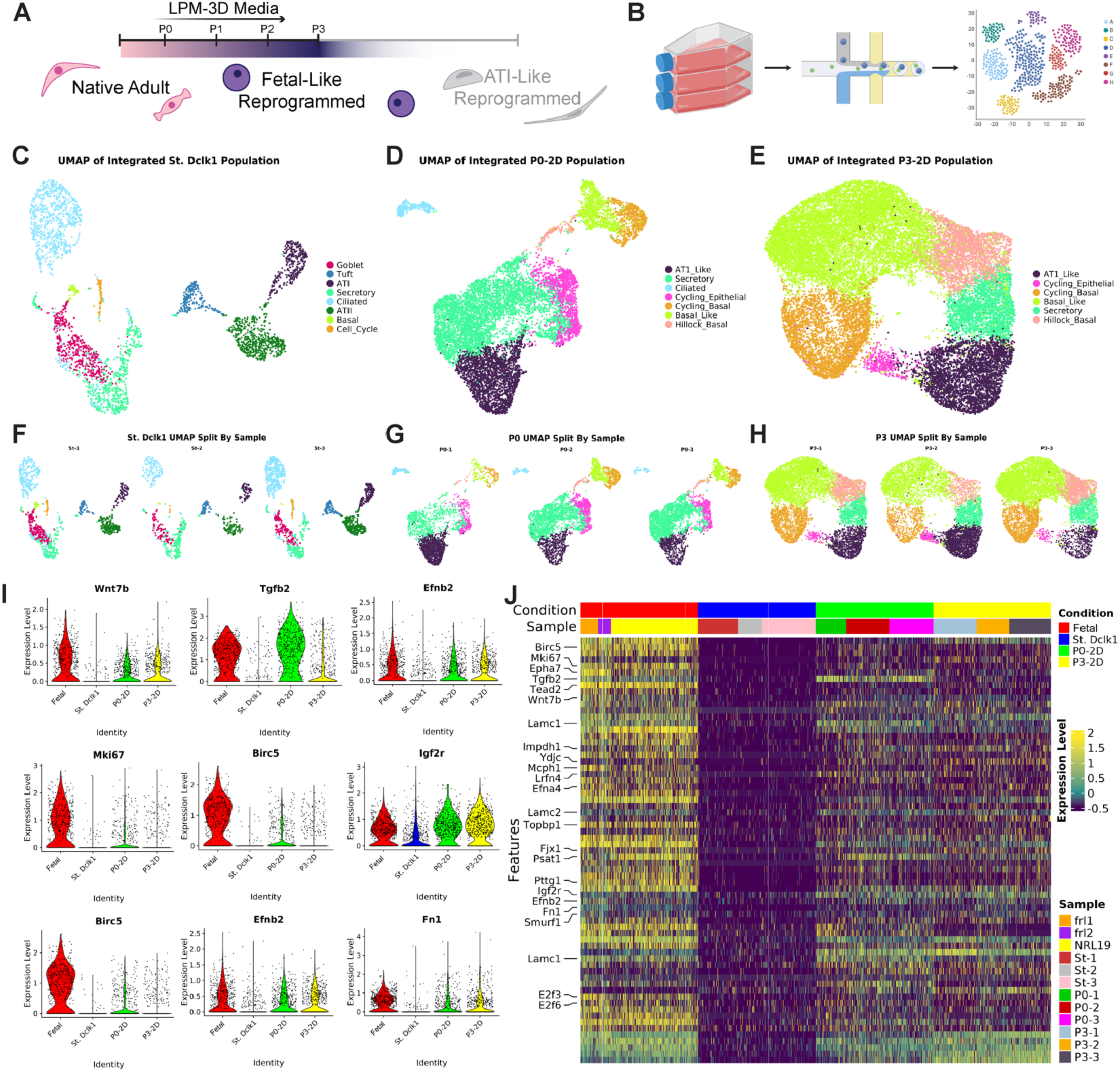
Characterization of reprogrammed epithelium. **A)** Schematic of cell state transition induced by LPM-3D. **B)** Schematic of 10x scRNA Sequencing Genomic Workflow. **C-E)** Condition-wise embeddings and clustering annotations following cleaning, iterative metadata construction, and integration. Analogous cell populations across conditions share the same color to highlight changes in cell number. **F-H)** Embeddings split by experimental replicate for each condition, respectively. Note the loss of cellular heterogeneity as populations progress from Starting to P0 to P3. **I)** Transcriptomic trends of interest. Fetal genes (*Tgfb2*, *Mki67*, *Tead2*, *Birc5*) that are downregulated in the adult dissociation become re-elevated in the 2D cultures. **J)** Heatmap depicting the scale of this change across many molecular families.

### Matrix culture induces epithelial production of inflammatory & regenerative cues

We next sought to explore how these reprogrammed cells would respond when seeded into decellularized lung and exposed to bare, acellular matrix. Following a protocol described in (*31*) and summarized in **Supplemental Figure 3**, we seeded 75 million P3 reprogrammed cells into the airways of decellularized rat lung matrices (n=3, biologic replicates) and cultured the constructs with 40 mL/min vascular perfusion and 4 bpm ventilation in LPM-3D media for 24 hours (**Figure 3A-B**). After 24 hours, the lungs were removed from the bioreactor and the lobes processed for analysis, including paraffin and frozen sectioning, bulk RNA isolation, and scRNAseq. **Supplemental Figure 4** shows global histology for all replicates. H&E staining showed living cells attached throughout the airways and alveoli, with higher cellular density and collective organization occurring at the adventitial and sub-pleural boundaries (**Figure 3A**). The cells displayed evidence of local proliferation while maintaining cuboidal morphology regardless of observed location in either alveoli or conducting airways (**Figure 3C-E**). Fluid flow modeling using data from culture (*32, 33*) showed no significant change in global tissue mechanics during the short culture time (**Figure 3F**). Immunohistochemistry of alveolar regions of the lung showed clear cellular expression of *Sox2*, *Sox9*, and *Tp63*, but did not show clear cellular expression of ATI-markers *Pdpn* (RTI-40), *Sema3e*, or *Vegfa*, suggesting that alveolar matrix and LPM-3D media combination was insufficient to induce full ATI-cell program expression and that the seeded cells were maintaining fetal-like character at 24 hours.

**Figure 3.**
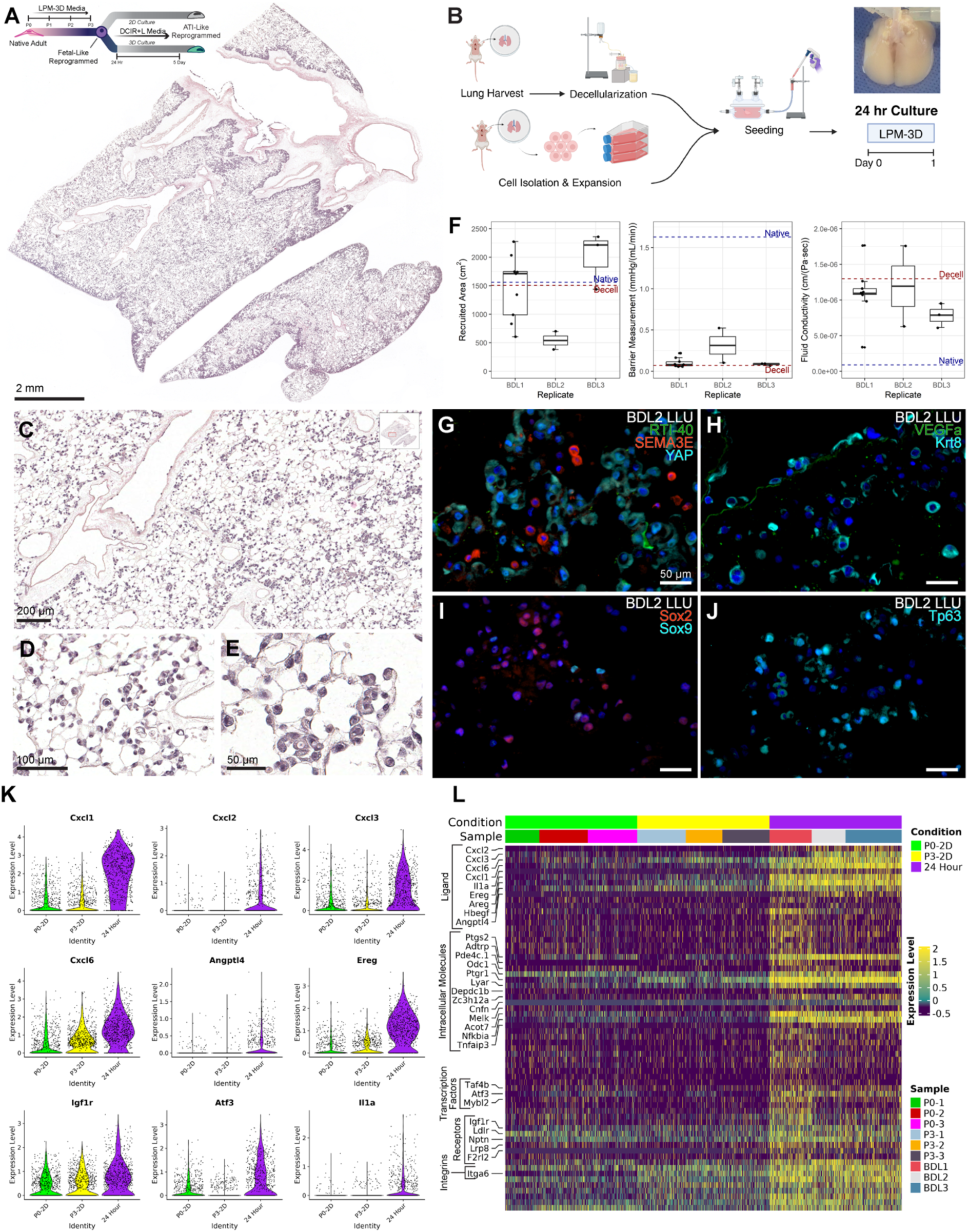
Epithelial cells cultured on bare matrix produce inflammatory & regenerative cues. **A)** Whole lobe cross section H&E showing excellent seeding coverage of reprogrammed epithelium into decellularized lung matrices. Scale bar: 2 mm. **B)** Schematic of experimental procedure. The cells are seeded into the airways of the lung and cultured in LPM-3D media for 24 hours. **C)** A higher magnification image demonstrating uniform distribution of cells exclusively in the airways, as desired. Scale bar: 200 µm. **D-E)** High-magnification H&E fields of view showing rounded cell morphology. Scale bars: 100 µm (D), 50 µm (E). **F)** Barrier modeling outputs for each replicate. **G-J)** Immunostaining of cells within the tissue at experimental endpoint. The cells are negative for *Pdpn*, *Sema3e*, and nuclear *YAP*, suggesting they have minimal ATI or ATI-transitional character. Conversely, they showed broad positivity for *Krt8*, *Sox2*, and *Tp63*, and patchy positivity for *Sox9*. Scale bars: 50 µm. **K)** Selected genes of interest showing significant elevation in the Engineered 24-Hour condition vs. P3 and P0 samples as controls. Epithelial cells increased their transcription of pro-inflammatory response genes (*Cxcl1*, *Cxcl2*, *Cxcl3*, *Cxcl6*, *Il1a*) concurrent with tissue repair genes (*Areg, Ereg, Hbegf, Angptl4, Atf3*). **L)** Heatmap demonstrating the ligands, intracellular molecules, transcription factors, receptors, and integrins upregulated in the 24-Hour condition, as compared to the reprogrammed epithelial cells cultured in 2D at passages 0 and 3.

To test this hypothesis, we performed scRNAseq of cells freshly isolated from these engineered tissues (n=3 biological replicates, 23,588 cells passing QC, see **Supplemental Figure 2A-C** for details), and compared the transcriptional profile to that of the P0 and P3 cultured cells. This allowed us to isolate the effect of bioreactor culture on reprogrammed cellular state. Upon transition to the 3D rat lung scaffold, we observed an unexpected increase in inflammatory response, denoted by transcriptional upregulation of the chemokine ligands and neutrophil attractants *Cxcl1*, *Cxcl2*, *Cxcl3*, and *Cxcl6*, as well as the pro-inflammatory cytokine *Il1a* (*34–36*) and the inflammatory mediator *Angptl4* (*37*) **(Figure 3K-L)**. We also noted elevated transcription of cyclooxegenase-2 (*Ptgs2*), an enzyme involved in inflammatory responses through prostaglandin generation, and activating transcription factor 3 (*Atf3*), a transcription factor which negatively regulates *Ptgs2* (*38*) and has recently been established as an important regulator of lung regeneration and repair (*39*). In addition to this inflammatory activation, we also observed significant upregulation of the epidermal growth factor family ligands epiregulin (*Ereg*), amphiregulin (*Areg*), and heparin-binding epidermal growth factor-like growth factor (*Hbegf*). Epiregulin is upregulated during wound healing, inflammation, and angiogenesis, and is upregulated in lung epithelium undergoing infection or compressive stress (*40*). Similarly, *Areg* is produced during inflammation and promotes tissue repair (*41*), and plays a role in bronchial epithelial regeneration following infection-induced injury (*42*). *Hbegf* also partakes in tissue repair and has been implicated in post-pneumonectomy alveolar regeneration (*43*) and post-injury tissue remodeling (*44*). These findings suggested that exposure to bare matrix in the bioreactor environment, in the absence of other cell types, was inducing epithelial inflammation and the transcription of tissue repair genes but was insufficient to promote transition to mature homeostatic states.

### Induction of ATI-cell programming in 2D

Within the alveolar regions of the engineered lungs, we observed occasional instances of cellular flattening, which lead us to hypothesize that we might be able to drive the reprogrammed cells directly to an ATI-cell state. To test this hypothesis, we compared the effect of culturing P4 cells in LPM-3D media vs. DCIR+L media adapted from Burgess et al. (*45*) (**Figure 4A**), a YAP-pathway-activating culture medium designed to induce ATII-to-ATI transition. **Figure 4B** outlines the composition of each media, including the Media Foundation which they share, and the media-specific components. Brightfield microscopy images were taken on Day 1, 3, and 5 of culture for both conditions grown in parallel and are shown in **Figure 4C**. Compared to the LPM-3D media cultures, the cells in the DCIR+L media condition did not proliferate as rapidly, and did not reach confluence until Day 5. By Day 1, the edges of the cells were not smooth as they had been in the LPM-3D condition, and many exhibited finger-like protrusions on edges that did not have contact with neighboring cells. By Day 3, the cells grown in the DCIR+L media begin to have an observable greater average cell surface area. By Day 5, most cells have a flat morphology with a large surface area, consistent with ATI-like morphology.

**Figure 4.**
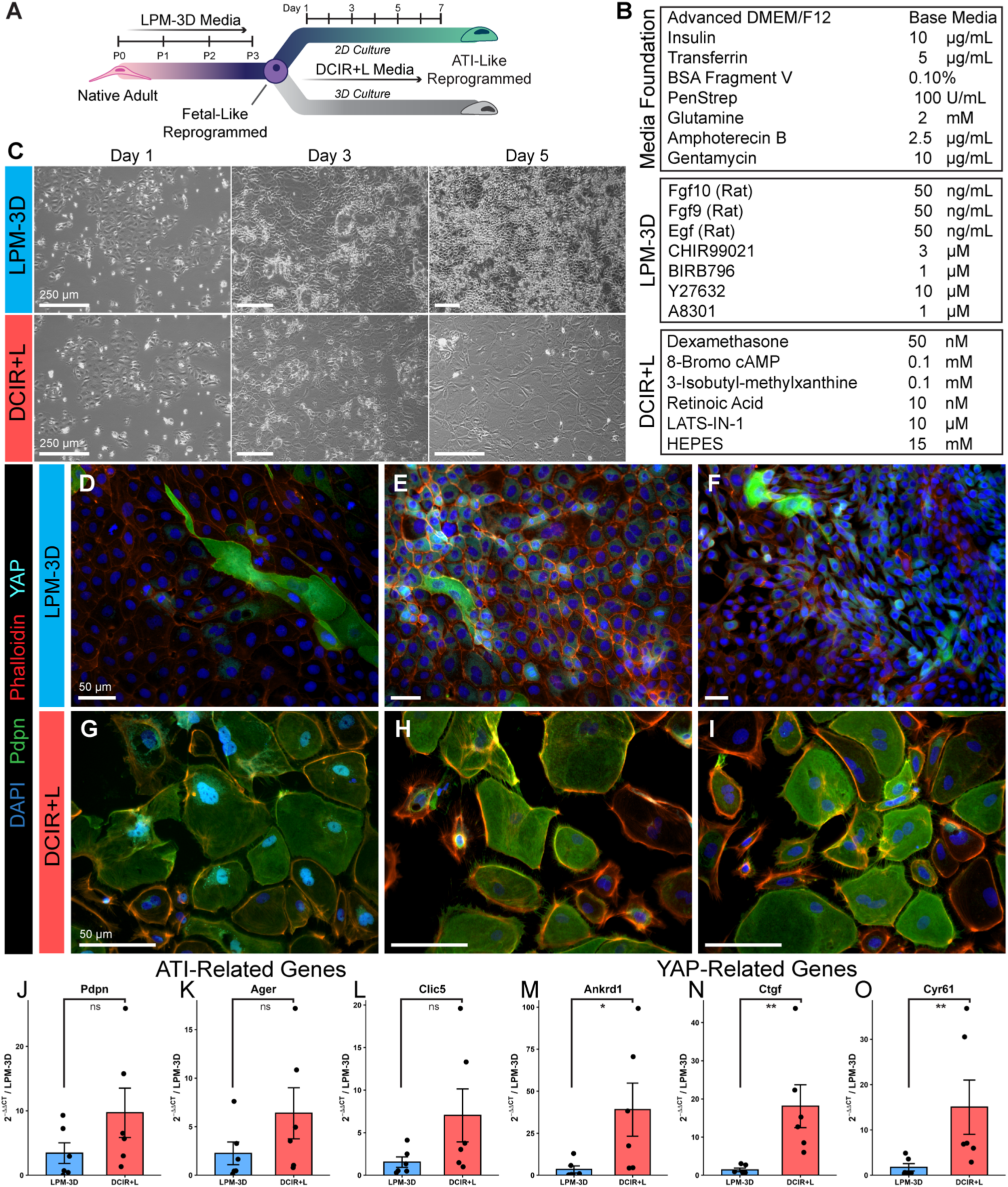
Induction of ATI-programming in 2D. **A)** Schematic of cell state transitions induced by this project, Part 2**. B)** Table of media compositions. Both media share the components of the media base, with additional components specific to the LPM-3D and DCIR+L media each. **C)** Brightfield microscopy comparing cell morphologies in LMP-3D and DCIR+L Media Conditions on Day 1, 3, and 5 in culture. Scale bars: 250 µm. Differences in cell morphology are apparent by Day 3, and obvious by Day 5. Namely, cells in the LPM-3D media are small, rounded, and grow to over-confluence, while cells in the DCIR+L media are larger in surface area, less compact, and generally do not overlap. **D-F)** LPM-3D Media Condition Fluorescent Microscopy, Frame 1. We observed no evidence of nuclear YAP and minimal Pdpn expression. Scale bars: 50 µm. **G-I)** DCIR+L media condition fluorescent microscopy. Scale bars: 50 µm. We observed patchy nuclear YAP translocation and widespread expression of *Pdpn*. Note the increased per-cell surface area, reminiscent of ATI cells. **J-O)** Quantitative RT-PCR for selected ATI and YAP-pathway related genes at Day 5 in each of the media conditions. Each dot represents a unique biological replicate and bars denote mean ± SEM. Statistical significance between LPM-3D and DCIR+L was assessed for each gene using a two-sided Wilcoxon rank-sum test on the ΔΔCt values; ns, not significant; *p < 0.05; **p < 0.01; ***p < 0.001.

Cells from both conditions were stained for *Pdpn*, *F-actin*, and *YAP*. Immunofluorescent images show *Pdpn* positivity in many of the large, flat cells cultured with the DCIR+L media, compared to few positive cells in the LPM-3D media condition (**Figure 4D-I)**. **Figure 4D-F** demonstrates how cells in LPM-3D have few *Pdpn*-positive cells, no nuclear-located YAP signal, and are small in size and densely distributed. **Figure 4G-I** demonstrate how the majority of cells in the ATI Differentiation Media are positive for Pdpn and have a large surface area with cells spread out across the plastic. **Figure 4G** shows nuclear-translocated YAP associated with the differentiation transition. The increased expression of ATI and YAP-related genes in the DCIR+L condition was consistent with RT-qPCR data performed on the cells. The data revealed a significant increase in the expression of *Pdpn*, *Ager*, *Clic5*, *Ankrd1*, *Ctgf*, and *Cyr61* among cells grown in DCIR+L media compared to the cells expanded in LPM-3D (**Figure 4J-O**), indicating transcriptional expression of ATI cell programming.

### ATI-program induction in whole lung culture

Armed with a way to induce ATI-directed transition in 2D, we sought to explore what the effect would be of culturing cells in 3D in DCIR+L within the bioreactor matrix environment. P3 cells were seeded into decellularized rat lung matrix and cultured in the LPM-3D media for 24 hours as before, but then switched to DCIR+L media on Day 1 (**Figure 5A-B**). Media exchanges were performed daily. Mechanical modeling showed insignificant changes in barrier properties or vascular recruitment during the 5-Day culture. At the end of the experiment, the lungs were removed from the bioreactor and processed in parallel for scRNAseq and paraffin and frozen sectioning. **Supplemental Figure 5** shows global histology and metabolic monitoring for all replicates.

**Figure 5.**
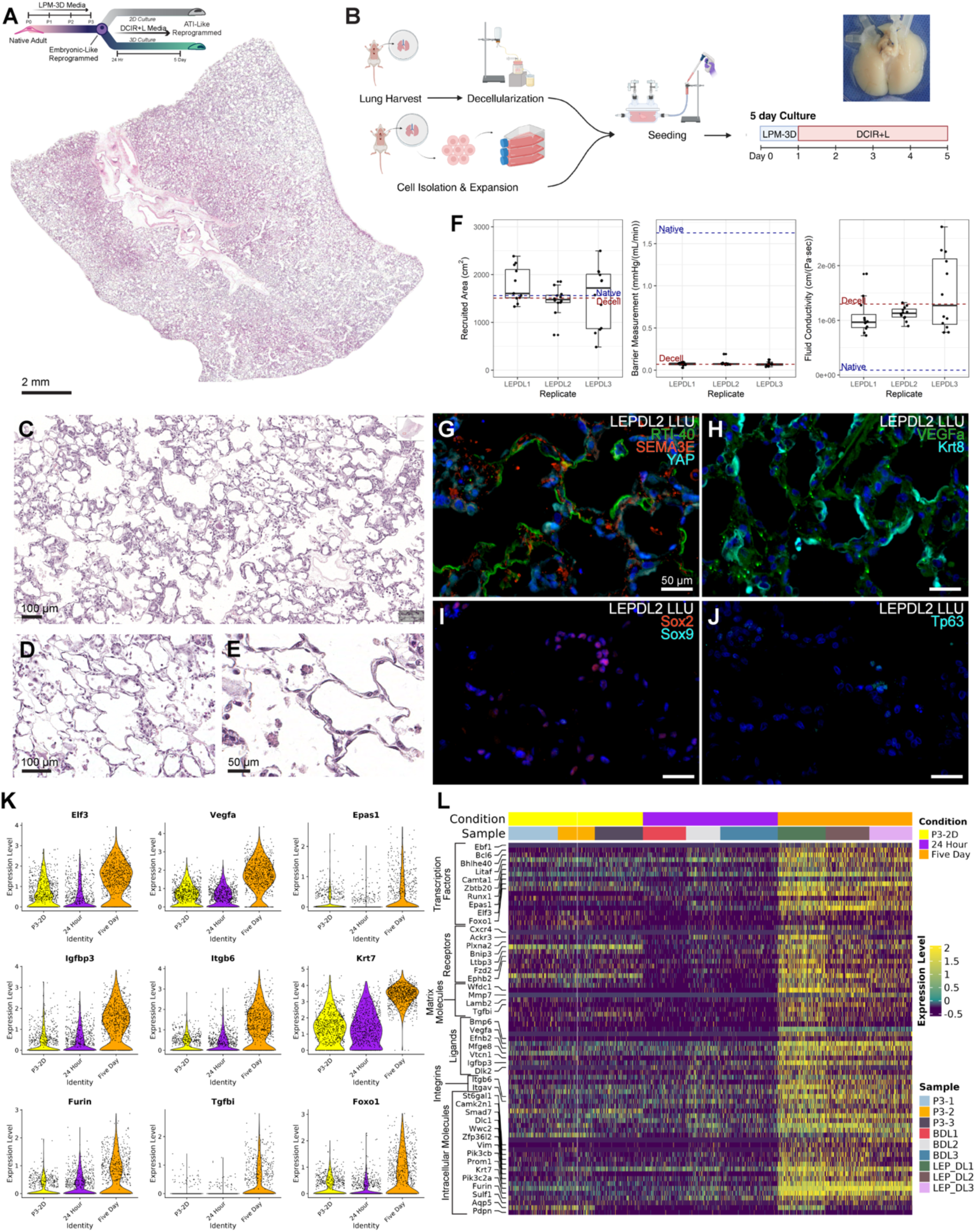
Induction of epithelial transition in 3-dimensional bioengineered lungs. **A)** Whole lobe cross section H&E showing homogenous seeding coverage. Scale bar: 2 mm. **B)** Experimental schematic for 5-day lung culture. Decellularized lungs were seeded with P3 reprogrammed epithelium, cultured in LPM-3D media for 24 hours, and then cultured in DCIR+L media for 4 additional days. **C-E)** Magnified H&E selections of alveolar territory showing flattening of cells against the alveolar walls. Scale bars: 100 µm (C-D), 50 µm (E). **F)** Mechanical data showing no significant change in the barrier capabilities of the tissue as compared to decellularized reference data. **G-J)** Immunostaining for selected markers. Scale bars: 50 µm. Note the upregulation of RTI-40 (*Pdpn*), *Sema3e*, and *Vegfa*, each of which mark ATI cells within native lungs. We observe a downregulation of transcription factors *Sox9*, *Sox2*, and *Tp63*, and an upregulation of the cytoskeletal protein *Krt8*. For layout allowing direct comparison between data from this condition and the 24-hour condition, see Supplemental Figure 2. **K)** Violin plots showing trends of interest. Induction of ATI programming in DCIR+L media causes a significant increase in ATI markers (*Vegfa*), genes associated with aberrant basaloid cells (*Itgb6, Krt7, Tgfbi)*, and several factors associated with cellular differentiation and remodeling (*Furin*, *Foxo1*, *Elf3).* **L)** Heatmap summarizing the epithelial state change induced by culture in DCIR+L. Note the broad involvement including transcription factors, receptors, matrix molecules, soluble ligands, integrins, and intracellular molecules. Transitional cells display several features of aberrant basaloid cells known from human disease, including *Efnb2*, *Mmp7, Lamb2, Itgb6, Vim, Krt7*.

Gross appearance and H&E staining confirmed uniform seeding and widespread viability (**Figure 5A**). Unique to the DCIR+L condition, we observed marked flattening of cells against the decellularized alveolar walls, in some cases approaching the <1µm cytoplasmic thinness seen in native ATI-cells (**Figure 5C-E**). IHC staining showed the cells to be positive for the ATI-cell markers *Pdpn*/RTI-40 and *Sema3e*. We also observed instances of both nuclear and cytoplasmic *YAP*, *Krt8*, and *Sox2*, while observing almost no expression of *Sox9* or *Tp63* (**Figure 5G-J**). These findings suggested that the cells were transitioning toward an ATI-like state while losing pluripotency and basal-like character in exchange.

To test this hypothesis, we analyzed scRNAseq data from the DCIR+L lungs (n=3 biologic replicates, 15,728 cells passing QC, see **Supplemental Figure 2D-F** for details), using both the P3 (2D) and 24-hour (3D) samples as controls. The 5-day condition showed clear differentiation of progenitors toward ATI programming. We observed increased transcription of *Krt7,* a cross-species marker of ATI cells, *Vegfa*, which is predominantly produced by ATI-cells in the lung, and *Sema3e*, which is an ATI-specific spatial-guidance molecule responsible for maintaining endothelial organization in the capillary-alveolar barrier. The transition toward ATI-character was associated with significantly decreased transcription of cellular-proliferation genes and significantly elevated transcription of cell-cycle arrest proteins, PI3 kinase genes, and tumor-suppressor genes, including the transcription factor *Elf3*. However, we found that the transitioning cells displayed many features that are not associated with ATI cells but instead are well known from the literature as biomarkers of transitionally-arrested ‘aberrant basaloid cells’ seen in IPF and COPD. After changing the media from LPM-3D to DCIR+L and incubation until day 5, we observed the upregulation of several markers associated with epithelial states now well-described in clinical lung pathology (**Figure 5K-L**)(*5, 46*). Our 5-Day population expressed integrin beta 6 (*Itgb6*), an integrin that is a long-established biomarker of pulmonary fibrosis and wound healing (*47, 48*), *Wnt7a,* a ligand expressed by both ATI-cells during homeostasis and aberrant basaloid cells during fibrotic remodeling (*49*), and ephrin type-B receptor 2 (*Ephb2*), a spatial guidance cue involved in postnatal alveolar development (*50, 51*) that is also a prominent marker of aberrant basaloid cells (*52*). We noted elevated transcription of vimentin (*Vim*), an intermediate filament upregulated in cells undergoing EMT (*53*) and matrix metalloproteinase 7 (*Mmp7*), suggesting cell-mediated matrix remodeling, and both of which mark aberrant basal cells in clinical data (*5*). ATI-marker gene *Aqp5* transcription was increased prominently in one of the three 24-Hour Conditions, while other ATI-associated markers, such as *Pdpn* showed only minimal transcription in 5-Day cells. These results collectively suggested incomplete transition toward an ATI-cell state, concurrent with adoption of an aberrant or arrested transitional phenotype similar to one seen in clinical lung pathology.

### Global observations and benchmarking of engineered cell states against native biologic archetypes

To formally test this hypothesis, we compared the cell states measured in this bioengineering context to cell states measured in native rat and human lungs. **Figure 6A-B** provides a visual overview of the cellular populations and tissues studied within this project. The fetal and adult tissues provided control data to contextualize the cell states observed within engineered 2D and 3D conditions, which, when proliferating in 2D culture, adopt morphology similar to that seen during native fetal tissue development. In 3D culture, we observed that the cells in the 24-hour lung culture still maintained fetal morphology, while the cells in the 5-day lung culture condition began to resemble the flattened cells lining the alveolar walls of the native adult lung without ever achieving the true physiologic thinness of fully differentiated ATI cells. This comparison suggests that the engineered transition from fetal-like to ATI-like adult cells in 2D and 3D culture may in some ways mirror the fetal to adult native transition and also suggests a structural linkage between cytoskeletal arrangement and transcriptional program activation in pulmonary epithelium.

**Figure 6.**
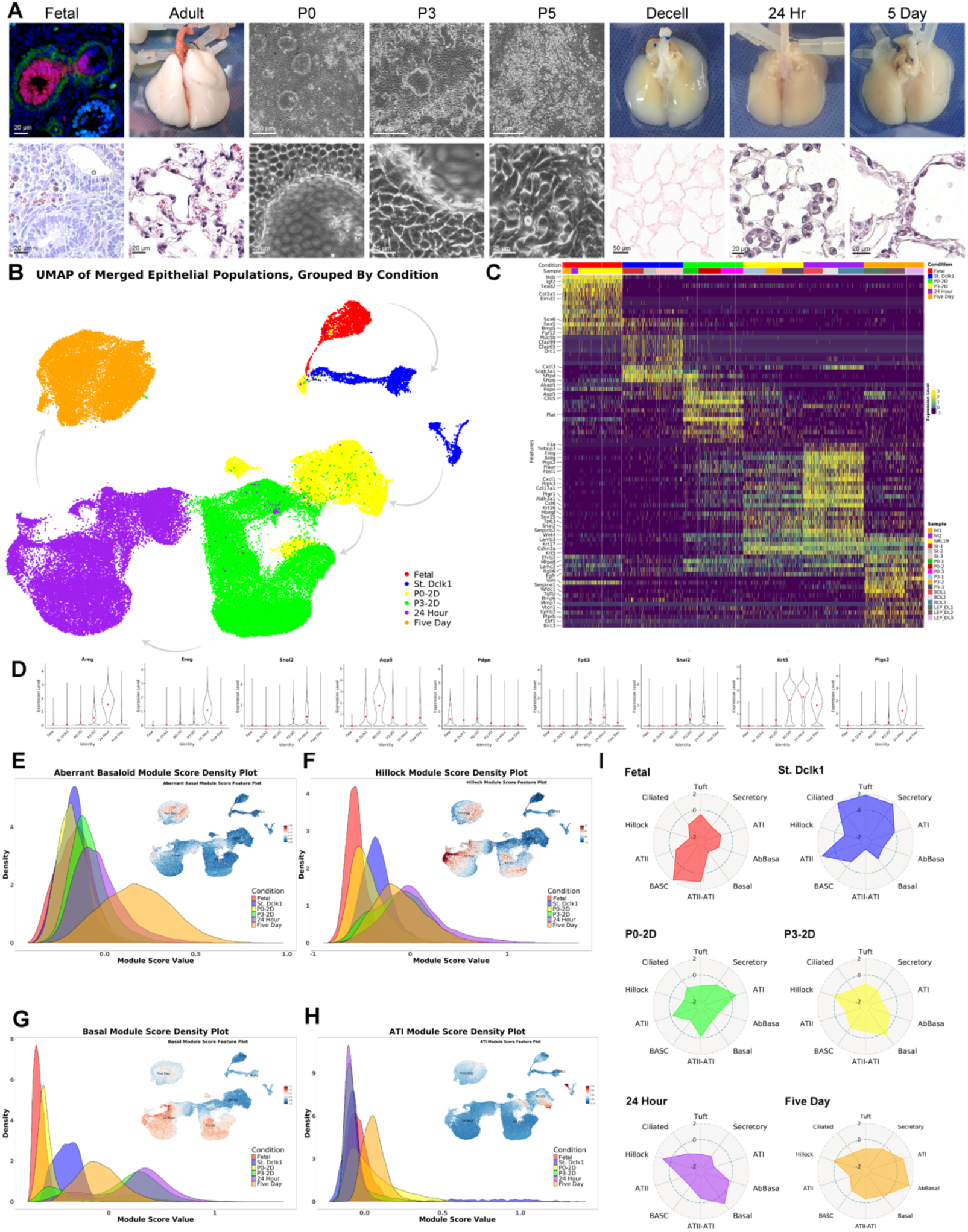
Engineered transitional cells in bioengineered lungs mirror aberrant basaloid cells seen in human clinical disease. **A)** Summary of the epithelial cell state landscapes profiled by this project. We have synthesized observations of fetal lung development, adult homeostasis, in vitro reprogramming, perfused matrix seeding, and induced transition. Scale bars provided with each respective image. **B)** Global UMAP of all cells, processed without integration. Grey arrows indicate progression from fetal → adult → passage 0 (LPM-3D) → passage 3 (LPM-3D) → 24 hour culture (matrix scaffold, LPM-3D) → 5 day culture (matrix scaffold, DCIR+L). **B)** Custom heatmap summarizing epithelial cell state distribution change across these 6 experimental arms. **D)** Condition-grouped violin plots detailing selected trends across experiments. Note the prominent upregulation of *Areg* and *Ereg* during the 24-hour LPM-3D culture which is downregulated via change to DCIR+L media, and the linkage between these regenerative genes and *Tp63* and *Krt5*, markers of basal cell character. Note also the inverse relationship with ATI markers *Pdpn* and *Aqp5*. **E-H)** Module scoring of global object using orthogonal gene queries derived from cell-state specific markers within native rat and native human data (see Methods). **E)** Transitional epithelial cells within the 5-day culture adopted aberrant basaloid character. **F)** Hillock cell character was adopted most prominently in the 24-hour LPM-3D condition. **G)** Basal cell scores (distinct from, and orthogonal to, Aberrant Basaloid scores) became downregulated upon culture in DCIR+L. **H)** ATI scores were highest in native, became downregulated during reprogramming culture, and are upregulated upon culture in DCIR+L. **I)** Radar plots summarizing cell state tendencies within each experimental condition. Fetal epithelium shows greatest similarity to adult stem cells and ATII-ATI transitional cells. Adult epithelium scores highly for well differentiated cells, including ATII, ATI, Ciliated, Tuft, and Secretory cells. In vitro 2D reprogramming through P3 drops scores for homeostatic cells while upregulating genes associated with both basal and hillock cells. Seeding onto perfused 3D matrix accentuates this trend, pushing the population even further from ATII and ATI cell differentiation while increasing both basal and hillock cell programming. Induction of epithelial transition induces both ATI cell programming and adoption of an aberrant basaloid phenotype known in human disease as a hallmark of alveolar tissue remodeling.

We performed a global analysis, encompassing all test and control data generated during these experiments, to characterize the major feature shifts associated with population-scale cell-state traversal. **Figure 6C-D** is a manually curated heatmap designed to visualize key global transcriptomic trends of interest. The switch to DCIR+L media induced a noticeable reversion of transcription of inflammatory mediators such as *Cxcl1*, *Cxcl2*, *Cxcl3*, *Cxcl6*, and *Il1a* in our 5-day condition, to levels nearing or below that seen in the 2D cultures. This decrease in inflammatory gene transcription coincided with a decrease in transcription of wound repair genes *Ereg*, *Areg*, and *Hbegf,* and a decrease in basal cell markers *Tp63*, *Krt5*, and *Krt14* (**Figure 6C-D**). This shift in cellular state was associated with a partial-but-incomplete differentiation towards ATI-cell character, with cells elevating their transcription of ATI-marker genes *Aqp5, Pdpn,* and *Clic5*. However, other genes which mark ATI cells *in vivo*, such as *Akap5* and *Ager,* show only minimal transcription that does not approach the levels measured in native ATI-cells. These trends collectively suggested that epithelial inflammation, regenerative ligand expression, and proliferation were linked to basal/basaloid/proximal cell character, while ATI-program activation is linked to reversion of all three of these genetic programs.

To directly test this hypothesis, we developed a benchmarking strategy to quantitatively score each cell within a native archetype space defined by known *in vivo* cellular reference states. Manually curating gene lists from the literature, we developed 10 fully orthogonal scoring axes (see **Supplemental Table 2**), with no gene symbol overlap, allowing us to place each cell within a 10-dimensional space reflecting degree of differentiation toward or away from each chosen phenotypic reference state (ATI, ATII-ATI, ATII, Aberrant Basaloid, Basal, BASC, Ciliated, Hillock, Secretory, & Tuft). We iteratively titrated each reference state gene list so that module scores, when applied to both native rat (*4*) and human (*5*) reference data, preferentially marked the targeted cell states while excluding as much as possible all other non-target cell states (see **Supplemental Figure 6**).

We then scored each individual engineered cell, from this entire project, within these 10 purpose-built and functionally validated orthogonal axes. Selected outputs from this method are shown in **Figure 6E-H**, with comprehensive outputs available in **Supplemental Figure 7**. The global analysis has been summarized in the form of 6 radar plots (**Figure 6I**), one for each condition, showing the globally scaled population-mean value for each orthogonal module score; overlaid radar plots can be seen in **Supplemental Figure 8A-B**. We note BASC and transitional ATII-ATI scores are dominant in the Fetal population. The native adult (‘St. Dclk1’) condition, as expected, scores highly for well-differentiated Tuft, Secretory, and Ciliated cell character. 2D culture in LPM-3D culture media reduced this differentiated character. In the Passage 3 and 24-Hour 3D conditions, we observed an increase in Basal and Hillock-like scores and a decrease in ATII and ATI-like character. During Five Day culture in DCIR+L, conversely, we observed an increase in ATI-like character, a decrease in Basal and Hillock-like character, and the highest Aberrant Basaloid scores measured across our entire study. Pairwise correlation analysis between all 10 module scores across all cells profiled (**Supplemental Figure 8C**) confirmed that ATI-directed differentiation was closely associated with the activation of aberrant basaloid programming and inversely correlated with basal and hillock gene transcription. Cell cycle scoring and cross plotting against module scores (selected plots shown in **Supplemental Figure 9**) confirmed that ATI-directed epithelial transition inversely correlated with cellular proliferation, and that the partially transitioned basaloid cells we created in this study displayed a transcriptomic signature consistent with cell-cycle arrest.

## Discussion

Replicable tissue bioengineering relies on our ability to deliberately navigate cellular state spaces for desired ends. In pulmonary biology, this task is complicated by a lack of high-quality quantitative data regarding what areas of state space are fundamentally available and/or unavailable in different experimental conditions and extracellular milieus. In particular, little is known about how native cellular state spaces (those biologically accessible and commonly observed *in vivo*) compared to engineered cellular state spaces (those available and either observed or inducible *in vitro*). This study sheds new, though certainly incomplete, light on this problem. We have shown that specific culture conditions remodel population heterogeneity, constrain epithelial cell programming, and can be predictably altered to drive cellular state transitions desirable for regenerative engineering and/or disease modeling.

The data in this paper makes clear that cellular markers which are highly specific to cell states seen commonly in native tissue contexts may not retain their meaning within an engineering context. The cell states that we observe in these experiments – within 2D culture, 24-hour matrix culture, and 5-day matrix culture, are not holistically identifiable as perfect orthologues of any cell states known to us from native lung biology in neither healthy nor diseased states, in neither human, mouse, rat, or pig biology. They represent cell states that are either specific to these precise experiments or have specific character associated with the regenerative processes that we have triggered therein. We suspect that the cellular states and state trajectories we have profiled here represent a small fraction of the state space that is truly available to living pulmonary epithelial cells when placed in different extracellular milieus.

The inflammatory patterns shown within our data are particularly interesting. Although inflammation is clinically thought of as a negative event associated with tissue destruction and dysfunction, here we observe inflammatory cues being produced by epithelial populations, in monoculture, within an engineered, sterile, non-mass-transfer-limited 3-dimensional tissue microenvironment. This is an unexpected finding, and one that we consider statistically robust given its clear and consistent presence across all replicates. Furthermore, our findings demonstrate a clear linkage between epithelial inflammatory programming, regenerative ligand transcription, and adoption of basal-cell-like character, as demonstrated by the cell state shift induced by 24-Hour bioreactor culture.

Our experiments in this area have allowed us to explore epithelial transition using a broad lens. By seeding reprogrammed epithelial cells onto decellularized, perfused extracellular lung scaffolds, and then inducing direct progenitor-to-ATI cell transition (without attempting to first induce and stabilize an intermediate ATII-cell state to mirror state trajectories found during *in vivo* epithelial homeostasis), we have unexpectedly driven cells into a *Krt17*^+^/*Itgb6*^+^ cellular phenotype that closely resembles transitionally-arrested aberrant basaloid epithelial cells seen in human lung pathology including IPF and COPD. This is an unexpected finding and one that was not deliberately engineered. It speaks to the ability of engineered ex vivo systems to recapitulate disease-like cellular states and opens many possible questions which might be explored and possibly answered using this system regarding how transitional cellular states are regulated and controlled.

Fundamentally, this paper shows that epithelial cells in the lung are highly plastic and are shaped by their extracellular milieu. Much recent literature has hinted that complete cellular differentiation may be inextricable from the differentiation of a *community* of cells — that proper tissue homeostasis may rely as much on the coordinated differentiation of local cellular niches as it does on the differentiation of the individual cells themselves. Our experiments here lend additional supportive, though far from conclusive, evidence to this broad theory. In engineered tissues lacking all non-epithelial lineages, we were able to induce ATI-cell programming but have been unable, thus far, to fully recapitulate ATI-cell differentiation, even when using culture media designed specifically for this purpose. We suspect that this may be because the epithelium requires properly structured information from neighboring alveolar niche cells to fully complete their transition into a stable ATI-cell state within the 3D culture environment. This hypothesis suggests opportunities for further experimentation in future studies leveraging multicellular engineered tissues.

Our study has several limitations. We do not know if the transitional arrest we observe is due to the lack of adjoining alveolar niche cells, improper media design or mechanical environment, or is simply a consequence of the short culture time. Future experiments should be performed to investigate each of these hypotheses. The experiments in this manuscript have used primary rat cells, as is common in the bioengineering field and for which the greatest amount of control and reference data exists, but future researchers may benefit from replicating these experiments using matched cells and scaffolds from other species, in particular mice (in which most in vivo genetic experiments have been performed), ferrets (which are an emerging model displaying physiologic lung architecture and terminal airways more closely mimetic of human) and/or human beings (which would provide the greatest evidence of clinical translational relevance). We have sought at every instance to perform highly controlled experiments and therefore maximize replicability, but cost and time limitations have forced us to limit ourselves to n=3 biologic replicates per experimental condition; additional biological replicates would strengthen the statistical power of this study and are likely warranted in the future. Finally, we have opted here, following much experimentation, to analyze our data without global cross-condition batch correction and/or integration. This is because our data displays a high-degree of (deliberately engineered) cross-condition variance in cellular state and phenotype, and we were not able to identify useful computational integration tools that could properly handle this variance without obscuring biologically relevant information and/or introducing additional unwanted artifacts. As computational tools develop that are better able to handle the broad cellular state-spaces encountered during joint analyses of engineered and native samples, we anticipate that our dataset may yield additional insights relevant both to fundamental pulmonary biology and bioengineering and might help to stress-test new computational tools seeking to bridge the gap between engineered and native tissue biology.

## Acknowledgements

The authors would like to thank Yale Pathology Tissue Services (YPTS), the Yale Center for Genomic Analysis (YCGA), and the Keck Biotechnology Research Laboratory. We would also like to thank Pavlina Baevova for her essential support.

## Data Availability

All sequencing data generated for this manuscript, including raw FASTQ files, extracted digital gene expression matrices, and processed R objects containing relevant meta-data have been made available at Gene Expression Omnibus under GEO accession **GSE315874**.

## Code Availability

Code written for all data analysis and figure craft related to this manuscript has been made publicly available on GitHub at https://github.com/RaredonLab/Mizoguchi2025.

## Author Contribu@ons

**Conceptualization:** SM, MSBR **Data curation:** VL, HK, SEE, NW **Formal analysis:** HK, NW **Funding acquisition:** TT, DS, TK, NK, MSBR **Investigation:** SM, VL, HK, SEE, NW, MTG, CD, CH, RR **Methodology:** SM, VL, HK, NW, SEE, TO, AG **Project administration:** SM, MSBR **Resources:** TK, MSBR **Software:** VL, HK, SEE, NW, MSBR **Supervision:** DS, TK, NK, MSBR **Validation:** SM, VL, HK, SEE **Visualization:** SM, VL, HK, SEE, NW, MSBR **Writing – original draft:** – SM, VL, HK, SEE, NW, MSBR **Writing – review & editing:** SEE, CD, DS, TO, AG, TT, TK, NK, MSBR

## Methods

### LPM-3D Media

The LPM-3D Fetal Expansion Media was adapted from the media described in Nichane et al., 2017(*21*), originally designed to grow *Sox9*^+^ fetal epithelial progenitors and adapted here for use reprogramming adult cells to an fetal state. For use with rat cells, rat-derived growth factors were substituted for the original mouse-derived factors. This media consists of advanced DMEM/F-12 (Gibco, 12634010), 5 µg/mL Heparin (Sigma-Aldrich, H3149-10KU), 10 ug/mL Insulin (Sigma-Aldrich, 91077C), 15 The LPM-3D Fetal Expansion Media was adapted from the media described in Nichane et al., 2017 (*21*), originally designed to grow *Sox9*+ fetal epithelial progenitors and adapted here for use reprogramming adult cells to an fetal state. For use with rat cells, rat-derived growth factors were substituted for the original mouse-derived factors. This media consists of advanced DMEM/F-12 (Gibco, 12634010), 5 µg/mL Heparin (Sigma-Aldrich, H3149-10KU), 10 ug/mL Insulin (Sigma-Aldrich, 91077C), 15 ug/mL Transferrin (Sigma-Aldrich, T8158), 0.1% BSA fragment V (Gemini Bio, 700-104P), 1% P/S (Gibco, 15140122), 1% amphotericin B (Cytiva, SV30078.01), 0.1% gentamicin (Gemini Bio, 400-100P-010), supplemented with 50 ng/mL Rat Egf (Peprotech, 400-25), 50 ng/mL Rat Fgf9 (Novus, NBP2-35196-5ug), 50 ng/mL Rat Fgf10 (Peprotech, 400-42), 3 µM CHIR99021 (Cayman Chemicals, 13122-10 mg), 1 µM BIRB796 (Cayman Chemicals, 10460-10 mg), 10 µM Y27632 (Cayman Chemicals, 10005583-50 mg), and 1 µM A8301 (Cayman Chemicals, 9001799-10 mg). Media components are shown in **Figure 2B**.

### DCIR+L Media

The DCIR+L ATI Differentiation Media was adapted from the formulation described by Burgess et al., 2024(*54*) with the addition of 10 nM retinoic acid (Sigma-Aldrich, R2625), to further promote ATI-like differentiation(*17, 55*). This media consists of advanced DMEM/F-12 (Gibco, 12634010), 15 mM HEPES (Corning, 25-060-CI), 10 ug/mL Insulin (Sigma-Aldrich, 91077C), 5 ug/mL transferrin (Sigma-Aldrich, T8158), 0.1% BSA fragment V (Gemini Bio, 700-104P), 1% P/S (Gibco, 15140122), 1% amphotericin B (Cytiva, SV30078.01), 0.1% gentamicin (Gemini Bio, 400-100P-010), supplemented with 50 nM dexamethasone (Sigma-Aldrich, D4902), 0.1 mM 8-bromoadenosine 3’,5’-cyclic monophosphate sodium salt (Sigma-Aldrich, B7880), 0.1 mM 3-isobutyl-1-methylxanthine (Sigma-Aldrich, I5879), and 10 µM LATS-IN-1 (Cayman Chemicals, 36623), which is a LATS-inhibitor that activates the YAP-associated Hippo pathway(*56*). The media composition is shown in **Figure 4B** (*56*). The media composition is shown in **Figure 4B**.

### Adult Rat Lung Harvest

Adult rat lungs were harvested from 230-250g male Sprague-Dawley rats, which were approximately 6-8-week-old. Ketamine (75 mg/kg) and Xylazine (5 mg/kg) working solution was injected intraperitoneally to anesthetize animals, followed by heparin (400 U/kg) to prevent blood clotting in the lungs. The thoracic cavity was accessed trans-diaphragmatically, and the trachea, lungs, and heart were exposed by removal of the anterior rib cage. The trachea and pulmonary artery (PA) were cannulated with Y-shaped 1/16-inch barbed fittings (Cole Parmer), and blood was cleared out by PA perfusion with PBS containing heparin (100 U/mL) and sodium nitroprusside (0.01 mg/mL). The lungs and heart were collected *en bloc* and transferred to a petri dish for further experimentation.

### Fetal Rat Lung Harvest

Fetal rat lungs were harvested at gestational stage E17, corresponding to the canalicular stage. Pregnant female Sprague-Dawley rats were anesthetized with inhaled isoflurane at 4% vol/vol and maintained at same condition. The rats were injected with sodium heparin (400 U/kg) intraperitoneally, followed by the administration of analgesia with subcutaneous injection of lidocaine (dose), meloxicam (1 mg/kg), and buprenorphine (EthiqaXR, 3.25 mg/kg). The abdominal cavity was accessed by a midline laparotomy incision, and the gravid uterus was exposed. Fetuses were extracted by cesarian section. The fetuses were secured to a working surface by pinning and the thoracic cavity was accessed trans-diaphragmatically, then the trachea, lung and heart were exposed by removing the anterior rib cage. The fetal left ventricle was opened with scissors, and the lung was perfused through the pulmonary artery (PA) with PBS containing 100 U/mL sodium heparin and 0.01 mg/mL sodium nitroprusside. The fetal lung and heart were extracted *en bloc* and placed on a petri dish for further experiment.

### Rat Lung Cell Isolation

A dissociation solution comprised of warmed Dulbecco’s modified Eagles’s medium (DMEM) containing elastase (3 U/mL), collagenase/dispase (1 mg/mL), deoxyribonuclease I (DNase I; 20 U/mL) was used to dissociate the rat lung to a single-cell suspension. Adult rat lungs harvested as previously described were perfused with dissociation buffer through PA, then inflated through the trachea with dissociation solution. Each lung lobe was cut at the hilum and collected and submerged in dissociation solution in a conical tube and incubated for 20 min in a water bath at 37°C with gentle rocking. After incubation, the tissue was gently passed through a wire mesh strainer using a weigh spatula, then the strainer was rinsed with DMEM containing 10% FBS, 1% penicillin/streptomycin (P/S), 1% amphotericin B and 0.1% gentamicin to collect remaining cells. The tissue solution was centrifuged for 5 min at 300 × g to pellet the cells which were then resuspended with ACK lysing buffer (Gibco, A1049201) at a 1:1 ratio of pellet volume and incubated for 120 sec at room temperature. PBS with 0.01% BSA was added to the cell suspension, and spun down for 5 min at 300 × g. The pellet was resuspended with PBS with 0.01% BSA and filtered through a 70 µm cell strainer. The cells were centrifuged again for 5 min at 300 × g, resuspended, then filtered twice through a 40 µm filter. The single cell suspension was counted, assessed for viability, then diluted to 1 × 10^7^ cells/mL for Dclk1 enrichment.

### Dclk1 Enrichment

Dclk1 antibody was conjugated with Dynabeads M-280 sheep anti-rabbit IgG (Invitrogen, 11203D) according to the manufacturer’s instructions. Cells were labeled with Dclk1 conjugated Dynabeads by incubating them together for 30 min at 4°C with gentle tilting and rotation. Tagged cells were magnetically isolated using the DynaMag-5 magnetic rack and rinsed with 0.01% BSA in PBS for 4 times then spun down for 5 min at 300 × g. Supernatant was discarded, and the pellet was resuspended in LPM-3D medium supplemented with 0.5 µg/mL phenol-red free Matrigel (Corning, 356237), then counted, assessed for viability with 0.4% Trypan Blue (Gibco, 15250061), and plated in a 6-well plate.

### Lung Epithelial Progenitor Cell Culture and Expansion

0.75-1 × 10^5^ cells were plated on 6-well plates after cell isolation and Dclk1 enrichment and cultured with LPM-3D supplemented with diluted growth factor reduced phenol-red-free Matrigel (5 µg/ml, Corning,356237). Cells were cultured in a CO2 incubator (37°C, 5% CO_2_), and media exchanges were performed the day after plating and every other day thereafter until the end of culture. P0 cells were cultured for 10 days, while cells P1 were cultured for 7 days before passaging. Passaged cells were either used for further experiments or resuspended in freezing medium (90% fetal bovine serum + 10% DMSO) and stored in -80°C overnight, then transferred into liquid nitrogen to be stored as reserves.

### AT1 Induction for LEPs

P3 LEPs were cultured in LPM-3D supplemented with growth factor reduced phenol-red-free Matrigel (Corning, 56237) at a concentration of 5 µg/mL until day 7 and passaged. The subsequent P4 LEPs were cultured in LPM-3D for 1 day, and the medium was replaced with DCIR+L media. Cells were cultured up to 4 days in DCIR+L, and media changes were performed every other day.

### Decellularization of Rat Lungs

Lungs for decellularization were harvested as previously described from 300-350 g male Sprague-Dawley rats, which were approximately 8-12-week-old. The lungs, trachea, and heart were extracted en bloc and placed on a petri dish for cannulation and transfer to the decellularization apparatus. The pulmonary vein (PV) was cannulated with a 1/8-inch Y-shaped barbed fittings (Cole Parmer). Decellularization of lungs was performed based on sodium deoxycholate (SDC)-based protocol as previously described (*57*). In summary, the lungs were perfused through the PA with antibiotic/antimycotic solutions (10% P/S, 4% Amphotericin B, 0.4% gentamicin in PBS with Ca^2+^ and Mg^2+^), then rinsed with 0.0035% Triton X-100 (Invitrogen, HFH10) in PBS with ions. After 30 min incubation with benzonase endonuclease solution (20 U/mL) in airway at room temperature, lungs were rinsed with 1M NaOH in PBS, followed by a 3-step concentration gradient SDC solution (0.01%, 0.05% and 0.1%). Next, lungs were perfused with benzonase endonuclease solution (20 U/mL) through the PA and incubated for 1h at RT, then rinsed with 0.5% Triton X-100 (Invitrogen, HFH10) in 0.5M EDTA containing PBS. Furthermore, lungs were thoroughly rinsed with 2L of PBS and perfused with antibiotics/antimycotics solution in PBS, followed by 48-hour incubation with slow PA perfusion at 8 mL/min, and finally stored at 4°C until use. Decellularized lungs were used for further bioengineering experiments within no more than 1 week.

### Bioreactor Apparatus

The bioreactor used was designed in-house and custom fabricated by Daryl Smith and Preston Smith at the Yale University Glass Shop. The chamber is made up of two parts: a glass basin with twelve lateral ports around the perimeter and a glass lid with six vertical ports of varied sizes. The bioreactor basin contains the media, tissue, and cells, and facilitates mass transfer internally while interfacing with external apparatus components. The bioreactor lid creates a closed system and has ports to facilitate mechanical conditioning and chemical monitoring. Typically, only two vertical ports of the lid are used. One is used to interface the closed bioreactor environment with a vacuum to apply constant negative pressure. The second one is used to interface the bioreactor with a syringe pump to apply cyclical changes to the vacuum, inducing a ventilation effect. Connections are made between the external and internal environments of the bioreactor using port connectors. All port connectors consist of a hollow screwcap fitted with a glass tube. The glass tube is fitted with a short piece of silicone tubing and a female luer-to-hose barb connector fitting. The design and use of port connectors allows us to dynamically manipulate the internal bioreactor environment without compromising sterility. There are three distinct types of port connectors that are used to accomplish distinct functions. The first is a Short Flat port connector which has a glass tube with a wide internal end and no fitting for simple measurements or draining. The second is a Mid Double port connector which has a glass tube with a narrow end fitted with silicone tubing for making connections within the bioreactor, including to the lung. A Mid Double port connector is used to make a Drop Line port connector which is fitted with perpendicular tubing to draw media from the bottom of the basin for perfusion or media exchange and sampling. There are six external flow circuits that are connected to the basin, including the Pulmonary Artery (PA), Pulmonary Vein (PV), and Trachea (T) which interact directly with the lung, and the Pleura, Hollow Fiber Cartridge (HFC) and Feed Line which only directly interact with the media. The complete bioreactor apparatus consists of the bioreactor lid and basin, Perfusion Circuits, Pressure Transducers, a Hollow Fiber Cartridge, a Breathing Column, a Feed Line, a PES Filter, a hybrid Pulse Dampener and Bubble Trap device, a Vacuum Pump, and a Syringe Pump. These elements work together to achieve sterility, mass transfer, physiologic control, and real-time culture monitoring.

### Whole Lung Bioengineering with Lung Epithelial Progenitors

Whole decellularized rat lungs were cultured in the above-described bioreactor platform with a continuous pressure/flow monitoring system. Briefly, acellular lung scaffolds were mounted in a sterile glass bioreactor specially designed for whole rat lung culture and were perfused with media in a CO_2_ incubator (37°C, 5% CO_2_) at a rate of 20 mL/min with a peristaltic pump. This Pre-Conditioning process before the introduction of cells ensures thorough infiltration of media into the lung construct and allows any air bubbles to pass as to not obstruct the Seeding of cells or perfusion of media to them during culture. For cell Seeding, 90-125 × 10^6^ lung epithelial progenitors (LEPs) suspended in 10 mL of media were introduced into the airways of the lung dECM by gravity and negative pleural pressure applied within the bioreactor (target of 10-15 mmHg transpulmonary pressure). After 45 min of static incubation in a CO_2_ incubator, perfusion of media through the PA began at 8 mL/min, then the rate of perfusion was increased in increments of +4 mL/min every 30 mins up to 20 mL/min. It was then increased in increments of +7 mL/min, every 60 min up to 40 mL/min. Ventilation was added at the PA ramp-up rate of 20 mL/min, starting at Δ1 mmHg transpulmonary pressure (P_tr_ – P_pl_) with a syringe pump, then increased in increments of +1 mmHg simultaneously with the perfusion rate ramp up to a final ventilation variation of Δ4 mmHg.

Six lungs were cultured in two different media and culture length conditions. Three lungs were cultured for 24 hours in LPM-3D media only, and another three lungs were cultured for 4 additional days in DCIR+L media. Close and continuous monitoring of the culture conditions from beginning to end is important for evaluating the state of the tissue and making informed decisions about tuning the conditions. Real-time data is collected by pressure transducers that are placed at key points in the bioreactor circuit to allow for monitoring of the internal pressure environment of the lung during culture. Media was also sampled every morning of culture to monitor the lactate, glucose, and pH levels.

Full-volume medium exchange was performed after 24 hours of culture for switching LPM-3D media with DCIR+L media. Half-volume media exchange was performed when media sampled with lactate less than 10 mM, while full-volume medium exchange was performed in a case of lactate greater than 10 mM. Lungs were removed from the bioreactor at Day 1 or Day 5, respectively, for each condition, and lobes were dissected for tissue dissociation for single-cell RNA sequencing and snap freezing each, and the remaining lobes were perfuse-fixed by gravity through PA with 4% paraformaldehyde for at least three hours then soaked overnight and used for histological analysis.

Throughout culture, the pressure transducers are continuously measuring the pressure of the PA, PV, Trachea, and pleura, while calculating the transpulmonary pressure which is the difference between the Trachea and the pleura pressures. Twice per day, barrier measurements were taken for the PA, PV, and Trachea. These measurements are performed by temporarily stopping flow in the circuit then restoring it to evaluate the instantaneous resistance to flow, per the exact protocol described in Engler et al. (*32*).

### Mechanical Data Processing

The measurements of the PA, PV, Trachea, and pleura pressure were processed using established MATLAB and R scripts from Raredon et al., 2021 to model and estimate the recruited area, hydraulic conductivity, and barrier pressures of each lung (*33*). The measurements of the PA, PV, Trachea, and pleura pressure were processed using established MATLAB and R scripts from Raredon et al., 2021 to model and estimate the recruited area, hydraulic conductivity, and barrier pressures of each lung (*33*).

Pressure transducers are included in the PA, PV, and Trachea circuits nearest to the lung and are strategically placed directly upstream from the flow from the lung for accurate intra-organ measurements. A pressure transducer is also attached to a Short Flat of the basin constituting the Pleura circuit for pseudo pleural measurements. LabChart was used to record continuous pressure measurement tracings of the PA, PV, Trachea, and Pleura, and to calculate a continuous transpulmonary pressure as the difference between readings of the Pleura from the Trachea. Twice per day, barrier measurements are taken for the PA, PV, and Trachea. These measurements are performed by temporarily stopping flow in each circuit, then restoring it to evaluate the instantaneous resistance to flow.

### Quantitative PCR (qRT-PCR)

To measure the relative gene expression for cell-type markers of interest in the isolated and 2D cultured epithelial populations and to validate immunohistochemistry, quantitative real-time polymerase chain reaction (qRT-PCR) was performed. Cell suspensions of cultured cells were homogenized in lysis Buffer RLT (Qiagen). Total RNA was isolated using the RNeasy Mini Kit (Qiagen) according to the manufacturer’s instructions. RNA was reverse transcribed using the iScript cDNA Synthesis Kit (Bio-Rad) according to the manufacturer’s instructions. cDNA was reverse transcribed in a thermocycler using the following procedure: 5 minutes at 25 °C, 20 minutes at 46°C, 1 min at 95°C, and then held at 4°C for 5 minutes. PCR reactions were run in triplicate using 2 μL of cDNA in a 26 μL final volume with iQ SYBR Green Supermix (Bio-Rad) and 0.5 μL of each forward and reverse primer (dependent on gene being tested; primer sequences listed below). qRT-PCR samples were processed using the CFX96 Real-Time PCR Detection System (Bio-Rad) using the following protocol: initial denaturation step of 4 min at 95 °C followed by 40 cycles of PCR for 15 s at 95 °C, 30 s at 60 °C, and 30 seconds at 72 °C. Average threshold cycle values (Ct) from PCR 19 reactions were normalized against β-actin expression for all samples and are reported as a fold change against β-actin using the 2-ΔΔCt processing method. Fold-change for gene of interest transcript levels between samples (ex: ‘A’ and ‘B’) were calculated as follows: 2-ΔΔCt where ΔCt = Ct (gene of interest) - Ct (β-actin) and ΔΔCt = ΔCt(A) - ΔCt(B).

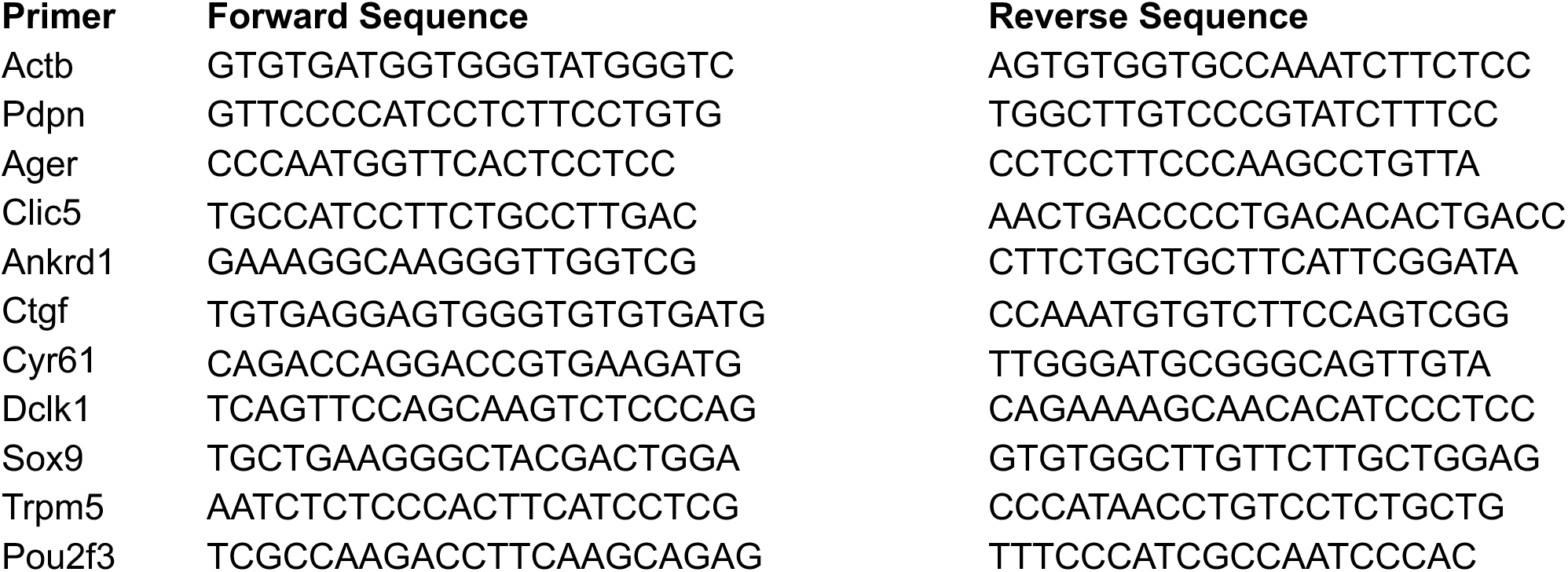

### BrighVield Microscopy

Brightfield microscopy images were captured every other day for all 2D cell cultures using Infinity Analyze software and the Carl Zeiss™ Axio Vert.A1 Inverted Microscope fitted with the Lumenera INFINITY2-1RM Monochrome CCD Camera. Areas of interest were captured with 4x, 10x, and 20x magnification objective lenses each time. The Auto-Exposure feature of the Infinity Analyze software was used at a setting of 50% to achieve consistent imaging of all samples.

### Cell & Tissue Fixing

Cells shown in situ were fixed to the plates on which they were grown. Cells processed via cytospin were collected in cell suspension and centrifuged at 1000 x g for 5 minutes onto slides using a Cytospin 4 Centrifuge (ThermoScientific), then fixed with 4% PFA.

At the end of each bioengineering culture, the lung is removed from the bioreactor and placed in a Petri dish for imaging of the gross appearance. For the purposes of the cultures performed for validating this method, the bronchiole is ligated at the hilum before one lobe is dissected for tissue dissociation and a single-cell RNA sequencing, another lobe is dissected for snap freezing and archive storage, and the remaining lobes are fixed by gravity perfusion through the PA with 4% PFA for at least three hours then soaked overnight on a rocker. The fixed lobes are dissected and sent for paraffin embedding and sectioning by Yale Pathology Tissue Services (YPTS). H&E, EVG, and trichrome histology staining are performed, and unstained tissue section slides are returned by the YPTS.

### Fluorescence Microscopy

IHC staining was performed for 2D-cultured cells and fixed tissues. Cell membranes were permeabilized using diluted Triton-X in PBS. Fixed and permeabilized cells were stabilized with Blocking Buffer by 1-hour incubation at room temperature, then incubated with primary antibodies (Table X) in Blocking Buffer at 4°C overnight. The following day, the cells were rinsed with PBS, then incubated with secondary antibodies in blocking buffer for 1 hour. After the secondary antibody incubation, the cells were rinsed, then mounted with PVA-DABCO and mounting glass.

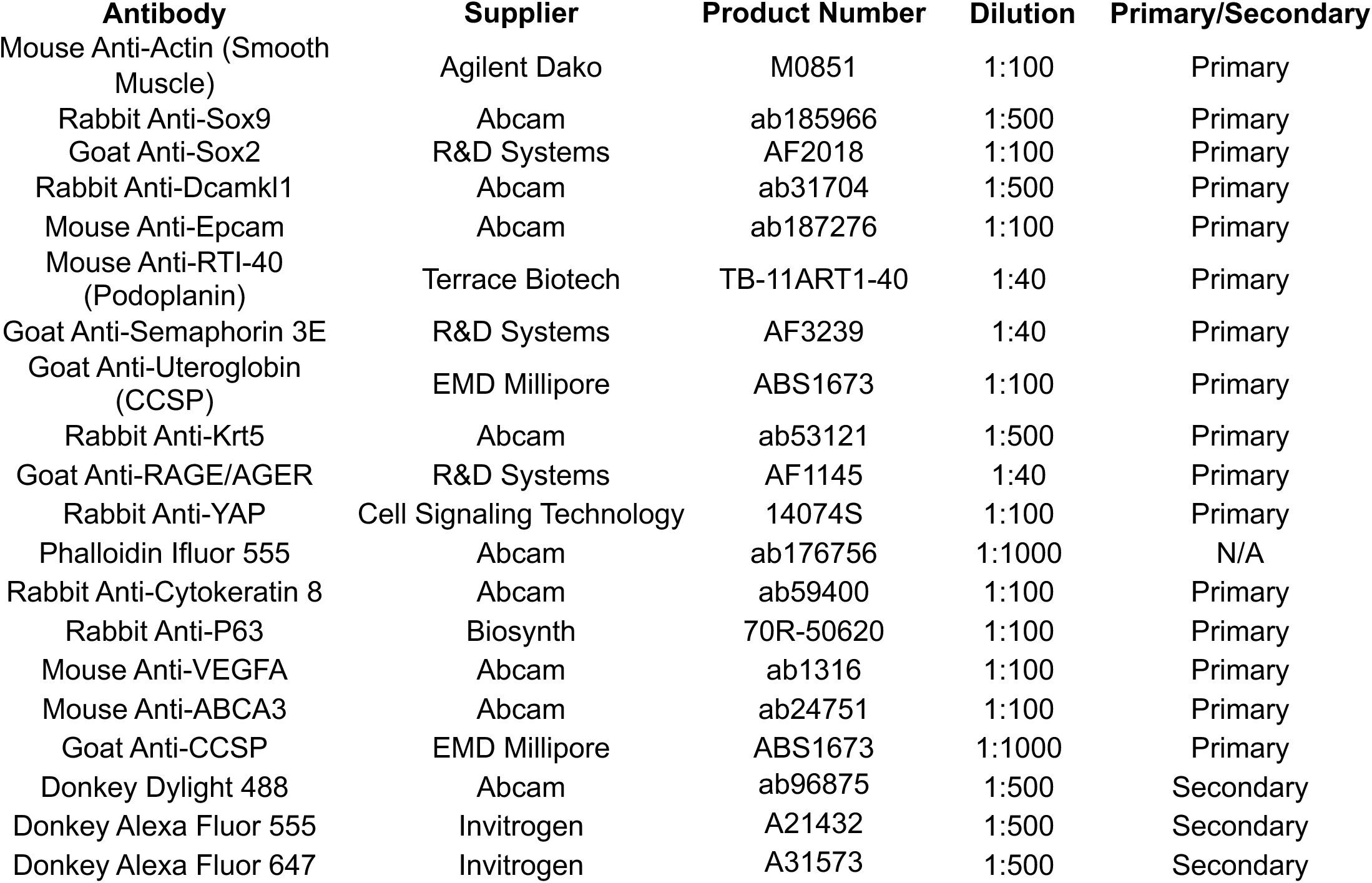

After staining, the cells were imaged using the EVOS FL Auto 2 fluorescent microscope to record single-frame, single-channel, and multi-channel single-frame and stitched frame images.

### Whole-Slide Digital Microscopy

H&E, EVG, and Trichrome IHC-stained tissue slides are sent to the Yale Pathology Tissue Services (YPTS) for digital microscopy, high-resolution, whole-slide scanning. Image files were analyzed using Aperio ImageScope and QuPath softwares.

### Image Quantification of Cytospin IHC Immunofluorescent Images

Cytospin slides were made using LEP cells at passages P1, P3, and P5. Two IHC assays were performed on three replicates of each condition. Assay 1 tested the cells for the presence of EpCAM, Ager, and Krt5. Assay 2 tested the cells for the presence of *Abca3*, *Sox2*, and *Sox9*. The immunofluorescent stained cells were imaged using the EVOS Fl Auto 2 fluorescent microscope to create stitched images of representative sections of the cytospin slides. The stitched images were processed using the Bioconductor package simpleSeg (*22*) The stitched images were processed using the Bioconductor package simpleSeg(*22*) to segment and characterize the cells on a single-cell level. The general workflow of image processing using this package involves loading the images of interest, reading them into R Studio using EBImage (*58*), converting them into a Large CytoImage (*58*), converting them into a Large CytoImage List (*59*), segmenting them according to defined parameters using the simpleSeg function, extracting the raw single-cell information, storing the information in a workable data frame, and processing and analyzing it. List (*59*), segmenting them according to defined parameters using the simpleSeg function, extracting the raw single-cell information, storing the information in a workable data frame, and processing and analyzing it. To ensure the most accurate segmentation was performed, segmented images were produced following iterations of variable parameters within the segmentation function. Only after satisfactory segmentation was achieved was the single-cell data extracted. The package provides each cell within each image an index ID, and extracts the source image title, x and y-coordinates, surface area in pixels, the major axis length in pixels, eccentricity, total number of segmented cells in the image, and the average pixel intensity of each channel within a cell.

### Single-Cell Library Preparation, Sequencing, & Alignment

Our single cell RNA seq analysis consists of six cell condition populations titled *Fetal*, *St. Dclk1*, *P0-2D*, *P3-2D*, *Engineered-24Hr*, and *Engineered-FiveDay*, with three replicates per condition. Single cell data for samples *St-1*, *St-2*, *St-3*, *P0-1*, *P0-2*, *P0-3*, *P3-1*, *P3-2*, *P3-3*, *BDL1*, *BDL2*, *BDL3*, *LEP_DL1*, *LEP_DL2*, and *LEP_DL3* were prepared using 10X Single Cell 3P v3.1 protocols. We aimed for a recovery of 10,000 cells per sample in our final library reads. Sequencing was done by the Yale Center for Genome Analysis (YCGA) on the Illumina NovaSeq 6000 platform. Fetal lung samples *frl1*, *frl2*, and *NRL19* were processed using the 10X Single Cell 3P RNA Seq kit, with final libraries depths sequenced to approximately 50,000 reads/cell. FastQ files were processed through CellRanger (*60*).

### Single-Cell Data Cleaning

Single cell data (raw, unfiltered digital gene expression matrices) were loaded into R using Seurat v5’s Load10X() (*61*). To retain as much information as possible when cleaning, we ran multiple “cycles” of variable feature generation, scaling, principal component analysis, and clustering to remove low-information cells. Our pipeline is a modified Seurat vignette for PBMC clustering.

For each sample, we filtered out cells based on factors such as *nCount_RNA* (total number of molecules in a cell) and *percent.mt* (percentage of a cell’s RNA that is mitochondrial). We then normalized our data, calculated the most variable features, and scaled our object. We generated and chose our desired principal components and created a highly clustered UMAP(*62*). Using visualization methods such as FeaturePlot() and ViolinPlot(), we further removed low-information clusters until we determined we were at risk of losing markers of interest with additional cleaning. We generated marker lists using the FindAllMarkers() function to double-check low-feature or high-percent.mt clusters for each sample. We created annotations for each sample based on existing literature, our marker lists, and past experimental findings. Following cleaning and annotation, we merged our 3D engineered samples into combined condition objects, then merged said objects together with previously generated 2D culture single cell objects, and fetal lung single cell samples. We subset and integrated each experimental condition via CCA integration, updated our annotations, and re-merged all into a unified Seurat object.

As an example, we detail the cleaning and annotation process for our BDL3 replicate. We created a count matrix from our data and used it to generate a Seurat object called BDL3 with 32,883 features (genes) across 2,440,680 samples (cells). We made a metadata slot, “percent.mt,” that informed us what percentage of a cell’s features are low-information mitochondrial genes. Next, we removed all cells that have an *nCount_RNA* below 500, and a percent.mt above 25. This removed many low-information cells, leaving us with 13,406 samples. We created and stored regular and log-scaled violin plots featuring *nCount_RNA*, *nFeature_RNA* (number of features in a cell), and percent.mt for BDL3 before and after thresholding. We normalized, calculated the 2000 most variable features, and scaled our data. Following this, we ran principal component analysis and chose principal components (PCs) 1 to 36 using dimensional reduction heatmaps and elbow plots to find the highest-variance PCs. Using the chosen PCs, we generated our first embedding and created a highly clustered UMAP. We determined which clusters have low *nFeature_RNA* and *nCount_RNA* with violin and feature plots. We also generated a marker list to look at the cluster feature expression levels, to confirm a cluster is low-information and can be removed. After sub-setting out low-information clusters, leaving us with 10,687 samples, we found variable features, scaled, and ran principal component analysis on our data again. We selected new PCs for embedding with dimensional reduction heat maps and elbow plots and created a second over clustered UMAP. We noted clusters with low *nFeature_RNA* for removal and checked their genes. Once we subset low-information clusters, leaving us with 10,128 samples, we found variable features, scaled, and ran principal component analysis again, selecting another group of PCs for embedding. We created a third over clustered heatmap, and examined the feature plots, violin plots, and marker lists for any low-information clusters. We finally determined our object to be completely cleaned, and re-clustered with a lower resolution. We created metadata slots for the sample name (“BLD3”), cell class (“Epithelial”), and the experimental condition (“Engineered-24Hr”) of our object, to use when BDL3 is merged with replicates and other experimental condition samples. We created a marker list using our simplified clusters and identified specifically expressed features, using existing literature as reference, and generating feature and violin plots for the markers. We changed the cluster numbers for our new labels based on our findings, saving the annotations in a new metadata identity. Lastly, we saved our cleaned and annotated BLD3 for future use.

### Single-Cell Data Analysis

We utilized Seurat’s FindMarkers() and intersect() function to create a set of reference lists of genes specific to an identity. We generated two overarching groups of reference lists, one with genes specific to experimental conditions, and the second with genes specific to each condition’s cell annotations. We also used FindMarkers() to construct marker lists of upregulated genes in the condition of interest for every permutation of condition comparisons in the merged Seurat object. For example, to generate upregulated marker lists for P0-2D, we made five Marker Lists comparing upregulated genes against Fetal, St. Dclk1, P3-2D, Engineered-24Hr, and Engineered-FiveDay. We created a variable titled *power* in all marker lists that was derived by multiplying the variables *avg_log2FC* and *ratio* (*ratio* itself was generated by dividing *pct.1* by *pct.2*). We filtered the top 2000 genes in order of descending *power* and created a character vector containing the row names (genes) of the marker list. We filtered out several extraneous genes to make our final feature vector.We then used intersect() to create gene lists tailored to our needs. For example, to create a list of genes specific to the Fetal condition, we ran multiple intersect() functions that combined the following: genes upregulated in Fetal over St. Dclk1, in Fetal over P0-2D, in Fetal over P3-2D, in Fetal Lung over Engineered-24Hr, and in Fetal Over Engineered-FiveDay. The result was a character vector that contained genes highly expressed in Fetal over all other conditions.

We displayed expressed genes by using an in-house wrapper function for Complex Heatmap (*63*). To adjust expressed gene ordering in our heatmaps, we created data frames containing our feature vector of interest, and corresponding variables from chosen marker lists. We changed the order of genes in this data frame as needed based on *pct.2* or *power*, with the aim of ordering genes that show more distinct expression levels between two conditions, before deriving a feature vector and creating the heatmap.

### Module Score Generation

We created ten sets of native features for module score calculation (ATI, ATII, ATII-ATI, BASC, Ciliated, Secretory, Tuft, Basal, Hillock, and Aberrant Basaloid). We used an existing native rat lung cell atlas to generate our native module score features. We first generated a feature list from our cell population of interest that was upregulated against the entire native rat lung cell atlas, then a second list with genes upregulated against the total epithelial native rat lung population, a third list with genes upregulated against the most spatially distant epithelial populations in the embedding, and finally a list of all genes upregulated against the most spatially proximal epithelial populations in the embedding. We then iteratively used R’s intersect() function to select a final list of highly specific features. After generating module genes, we used Seurat’s AddModuleScore() function to calculate the module scores for each condition. To generate our Aberrant Basaloid set of module genes, we used a human IPF atlas to find genes specific to the human aberrant basaloid population, then found murine orthologs to calculate the final module score. We generated FeaturePlots of both individual conditions and the global merged Seurat object with the module scores. We created a radar plot based on the mean values of each module using the ricardo-bion/ggradar package. For an accurate representation of score changes across each condition, we used the scale() function on each group of module condition scores. For visual clarity, we split the plot by condition to highlight changes in mean score.

## Bibliography

1. S. M. Rafelski, J. A. Theriot, Establishing a conceptual framework for holistic cell states and state transitions. Cell 187, 2633–2651 (2024).

2. B. Lim, K. Domsch, M. Mall, I. Lohmann, Canalizing cell fate by transcriptional repression. Molecular Systems Biology 20, 144-161-161 (2024).

3. M. Adler et al., Emergence of division of labor in tissues through cell interactions and spatial cues. Cell Reports 42, (2023).

4. M. S. B. Raredon et al., Single-cell connectomic analysis of adult mammalian lungs. Science Advances 5, eaaw3851 (2019).

5. T. S. Adams et al., Single-cell RNA-seq reveals ectopic and aberrant lung-resident cell populations in idiopathic pulmonary fibrosis. Science Advances 6, eaba1983 (2020).

6. M. Sauler et al., Characterization of the COPD alveolar niche using single-cell RNA sequencing. Nature Communications 13, 1–17 (2022).

7. N. J. Lang et al., Ex vivo tissue perturbations coupled to single-cell RNA-seq reveal multilineage cell circuit dynamics in human lung fibrogenesis. Science translational medicine 15, eadh0908 (2023).

8. A. Vannan et al., Spatial transcriptomics identifies molecular niche dysregulation associated with distal lung remodeling in pulmonary fibrosis. Nature Genetics 57, 647–658 (2025).

9. A. T. Abdallah, M. Peitz, A. Konermann, Revealing Genetic Dynamics: scRNA-seq Unravels Modifications in Human PDL Cells across In Vivo and In Vitro Environments. International Journal of Molecular Sciences 25, 4731 (2024).

10. M. L. Adelus et al., Single-cell ‘omic profiles of human aortic endothelial cells in vitro and human atherosclerotic lesions ex vivo reveal heterogeneity of endothelial subtype and response to activating perturbations. eLife 12, RP91729 (2024).

11. K. D. Alysandratos et al., Culture impact on the transcriptomic programs of primary and iPSC-derived human alveolar type 2 cells. JCI Insight 8, (2023).

12. C. Bock et al., The Organoid Cell Atlas. Nature Biotechnology 39, 13–17 (2021).

13. K. B. Jensen, M. H. Little, Organoids are not organs: Sources of variation and misinformation in organoid biology. Stem Cell Reports 18, 1255–1270 (2023).

14. B. E. Mead et al., Harnessing single-cell genomics to improve the physiological fidelity of organoid-derived cell types. BMC Biology 16, 62 (2018).

15. R. Gupta et al., Comparing in vitro human liver models to in vivo human liver using RNA-Seq. Arch Toxicol 95, 573–589 (2021).

16. Y. A. Kharaz, S. R. Tew, M. Peffers, E. G. Canty-Laird, E. Comerford, Proteomic differences between native and tissue-engineered tendon and ligament. Proteomics 16, 1547–1556 (2016).

17. K. L. Leiby et al., Rational engineering of lung alveolar epithelium. npj Regenerative Medicine 8, 22 (2023).

18. J. G. Camp, D. Wollny, B. Treutlein, Single-cell genomics to guide human stem cell and tissue engineering. Nature methods 15, 661–667 (2018).

19. Samantha A. Morris et al., Dissecting Engineered Cell Types and Enhancing Cell Fate Conversion via CellNet. Cell 158, 889–902 (2014).

20. A. H. Radley et al., Assessment of engineered cells using CellNet and RNA-seq. Nature Protocols 12, 1089–1102 (2017).

21. M. Nichane et al., Isolation and 3D expansion of multipotent Sox9+ mouse lung progenitors. Nature methods 14, 1205 (2017).

22. N. C. a. A. N. a. E. Patrick, simpleSeg: A package to perform simple cell segmentation. (2023).

23. J. Rajagopal et al., Wnt7b stimulates embryonic lung growth by coordinately increasing the replication of epithelium and mesenchyme. Development 135, 1625–1634 (2008).

24. L. P. Sanford et al., TGFbeta2 knockout mice have multiple developmental defects that are non-overlapping with other TGFbeta knockout phenotypes. Development 124, 2659–2670 (1997).

25. K. M. Bennett et al., Ephrin-B2 reverse signaling increases alpha5beta1 integrin-mediated fibronectin deposition and reduces distal lung compliance. Am J Respir Cell Mol Biol 49, 680–687 (2013).

26. N. Smyth et al., Absence of basement membranes after targeting the LAMC1 gene results in embryonic lethality due to failure of endoderm differentiation. J Cell Biol 144, 151–160 (1999).

27. J. Gerdes, U. Schwab, H. Lemke, H. Stein, Production of a mouse monoclonal antibody reactive with a human nuclear antigen associated with cell proliferation. Int J Cancer 31, 13–20 (1983).

28. G. Ambrosini, C. Adida, D. C. Altieri, A novel anti-apoptosis gene, survivin, expressed in cancer and lymphoma. Nat Med 3, 917–921 (1997).

29. M. Sokka, S. Parkkinen, H. Pospiech, J. E. Syvaoja, Function of TopBP1 in genome stability. Subcell Biochem 50, 119–141 (2010).

30. M. Makiniemi et al., BRCT domain-containing protein TopBP1 functions in DNA replication and damage response. J Biol Chem 276, 30399–30406 (2001).

31. A. M. Greaney et al., Engineered Whole Lungs for Tissue Biology. bioRxiv, 2024.2010. 2002.616240 (2024).

32. A. J. Engler et al., Non-invasive and real-time measurement of microvascular barrier in intact lungs. Biomaterials 217, 119313 (2019).

33. M. S. B. Raredon, A. J. Engler, Y. Yuan, A. M. Greaney, L. E. Niklason, Microvascular fluid flow in ex vivo and engineered lungs. Journal of Applied Physiology 131, 1444–1459 (2021).

34. C. Zhou, Y. Gao, P. Ding, T. Wu, G. Ji, The role of CXCL family members in different diseases. Cell Death Discov 9, 212 (2023).

35. E. T. Osei et al., Interleukin-1alpha drives the dysfunctional cross-talk of the airway epithelium and lung fibroblasts in COPD. Eur Respir J 48, 359–369 (2016).

36. A. Malik, T. D. Kanneganti, Function and regulation of IL-1alpha in inflammatory diseases and cancer. Immunol Rev 281, 124–137 (2018).

37. L. Guo et al., Role of Angptl4 in vascular permeability and inflammation. Inflamm Res 63, 13–22 (2014).

38. J. Hellmann et al., Atf3 negatively regulates Ptgs2/Cox2 expression during acute inflammation. Prostaglandins Other Lipid Mediat 116-117, 49–56 (2015).

39. T. K. Niethamer et al., Atf3 defines a population of pulmonary endothelial cells essential for lung regeneration. Elife 12, e83835 (2023).

40. D. J. Riese, 2nd, R. L. Cullum, Epiregulin: roles in normal physiology and cancer. Semin Cell Dev Biol 28, 49–56 (2014).

41. C. Berasain, M. A. Avila, Amphiregulin. Semin Cell Dev Biol 28, 31–41 (2014).

42. D. M. W. Zaiss, W. C. Gause, L. C. Osborne, D. Artis, Emerging functions of amphiregulin in orchestrating immunity, inflammation, and tissue repair. Immunity 42, 216–226 (2015).

43. D. T. Dao, L. Anez-Bustillos, R. M. Adam, M. Puder, D. R. Bielenberg, Heparin-Binding Epidermal Growth Factor-Like Growth Factor as a Critical Mediator of Tissue Repair and Regeneration. Am J Pathol 188, 2446–2456 (2018).

44. J. Ito et al., Wound-induced TGF-β1 and TGF-β2 enhance airway epithelial repair via HB-EGF and TGF-α. Biochem Biophys Res Commun 412, 109–114 (2011).

45. C. L. Burgess et al., Generation of human alveolar epithelial type I cells from pluripotent stem cells. Cell stem cell 31, 657–675.e658 (2024).

46. T. S. Adams, et al., Alveolar epithelial cell plasticity and injury memory in human pulmonary fibrosis. Biorxiv (Cold Spring Harbor Laboratory), (2025).

47. A. Meecham, J. F. Marshall, The ITGB6 gene: its role in experimental and clinical biology. Gene X 5, 100023 (2020).

48. A. L. Tatler et al., Amplification of TGFβ Induced ITGB6 Gene Transcription May Promote Pulmonary Fibrosis. PLoS One 11, e0158047 (2016).

49. G. Huang et al., Basal Cell-derived WNT7A Promotes Fibrogenesis at the Fibrotic Niche in Idiopathic Pulmonary Fibrosis. Am J Respir Cell Mol Biol 68, 302–313 (2023).

50. A. Vadivel et al., Critical role of the axonal guidance cue EphrinB2 in lung growth, angiogenesis, and repair. Am J Respir Crit Care Med 185, 564–574 (2012).

51. G. A. Wilkinson, J. C. Schittny, D. P. Reinhardt, R. Klein, Role for ephrinB2 in postnatal lung alveolar development and elastic matrix integrity. Developmental dynamics: an official publication of the American Association of Anatomists 237, 2220–2234 (2008).

52. C. H. Mayr et al., Spatial transcriptomic characterization of pathologic niches in IPF. Science Advances 10, eadl5473.

53. J. Ivaska, Vimentin: Central hub in EMT induction? Small GTPases 2, 51–53 (2011).

54. C. L. Burgess et al., Generation of human alveolar epithelial type I cells from pluripotent stem cells. Cell Stem Cell 31, 657–675.e658 (2024).

55. R.-w. Gao et al., Retinoic acid promotes primary fetal alveolar epithelial type II cell proliferation and differentiation to alveolar epithelial type I cells. In Vitro Cellular & Developmental Biology - Animal 51, 479–487 (2015).

56. H. Fujimori et al., LATS Inhibitor Protects 6-OHDA Induced Neuronal Cell Death In Vitro and In Vivo. BPB Reports 6, 144–149 (2023).

57. E. A. Calle et al., Targeted proteomics effectively quantifies differences between native lung and detergent-decellularized lung extracellular matrices. Acta Biomater 46, 91–100 (2016).

58. G. Pau, F. Fuchs, O. Sklyar, M. Boutros, W. Huber, EBImage—an R package for image processing with applications to cellular phenotypes. Bioinformatics 26, 979–981 (2010).

59. N. Eling, N. Damond, T. Hoch, B. Bodenmiller, cytomapper: an R/Bioconductor package for visualization of highly multiplexed imaging data. Bioinformatics 36, 5706–5708 (2021).

60. G. X. Y. Zheng et al., Massively parallel digital transcriptional profiling of single cells. Nature Communications 8, 14049 (2017).

61. Y. Hao et al., Dictionary learning for integrative, multimodal and scalable single-cell analysis. Nature Biotechnology 42, 293–304 (2024).

62. L. McInnes, J. Healy, J. Melville, Umap: Uniform manifold approximation and projection for dimension reduction. *arXiv preprint arXiv:1802.03426*, (2018).

63. Z. Gu, R. Eils, M. Schlesner, Complex heatmaps reveal patterns and correlations in multidimensional genomic data. Bioinformatics 32, 2847–2849 (2016).

