## Supplemental Figures for "Epithelial Reprogramming and Transition during Pulmonary Bioengineering"

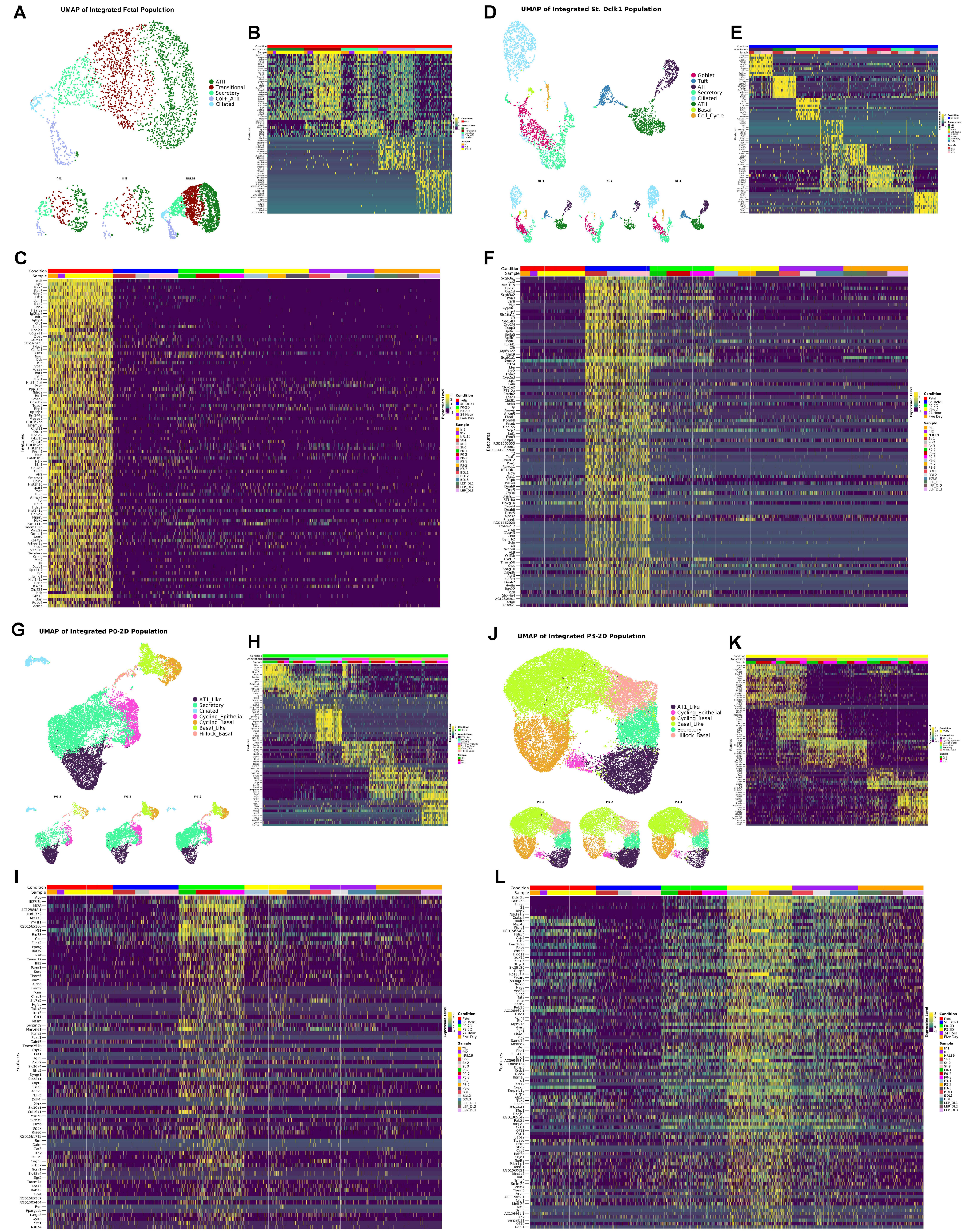


**Supplemental Figure 1.** Fetal, adult, and 2D cultured (passages 0 and 3) lung epithelium profiled by scRNAseq. A) Cross-sample UMAP embedding for fetal lung showing annotated cell states and the global embedding split by replicate. B) Heatmap showing genes marking each cell type cluster within the fetal lung epithelium object. C) Heatmap showing genes marking fetal lung epithelial against all other conditions in this study. D) Cross-sample UMAP embedding for adult lung epithelium showing annotated cell states and representation in each sample. E) Heatmap showing genes marking each cell type cluster within the adult lung epithelium object. F) Heatmap showing genes marking adult lung epithelial against all other conditions in this study. G) Cross-sample UMAP embedding for reprogrammed lung epithelium (P0) showing annotated cell states and the global embedding split by sample (replicate). H) Heatmap showing genes marking each cell type cluster within the reprogrammed lung epithelial (P0). I) Heatmap showing genes marking the reprogrammed lung epithelium (P0) against all other conditions in this study. J) Cross-sample UMAP embedding for reprogrammed lung epithelium (P3) showing annotated cell states and the cross-sample embedding split by sample (replicate). K) Heatmap showing genes marking each cell type cluster within the reprogrammed lung epithelium (P3) object. L) Heatmap showing genes marking the reprogrammed lung epithelial (P3) against all other conditions in this study.

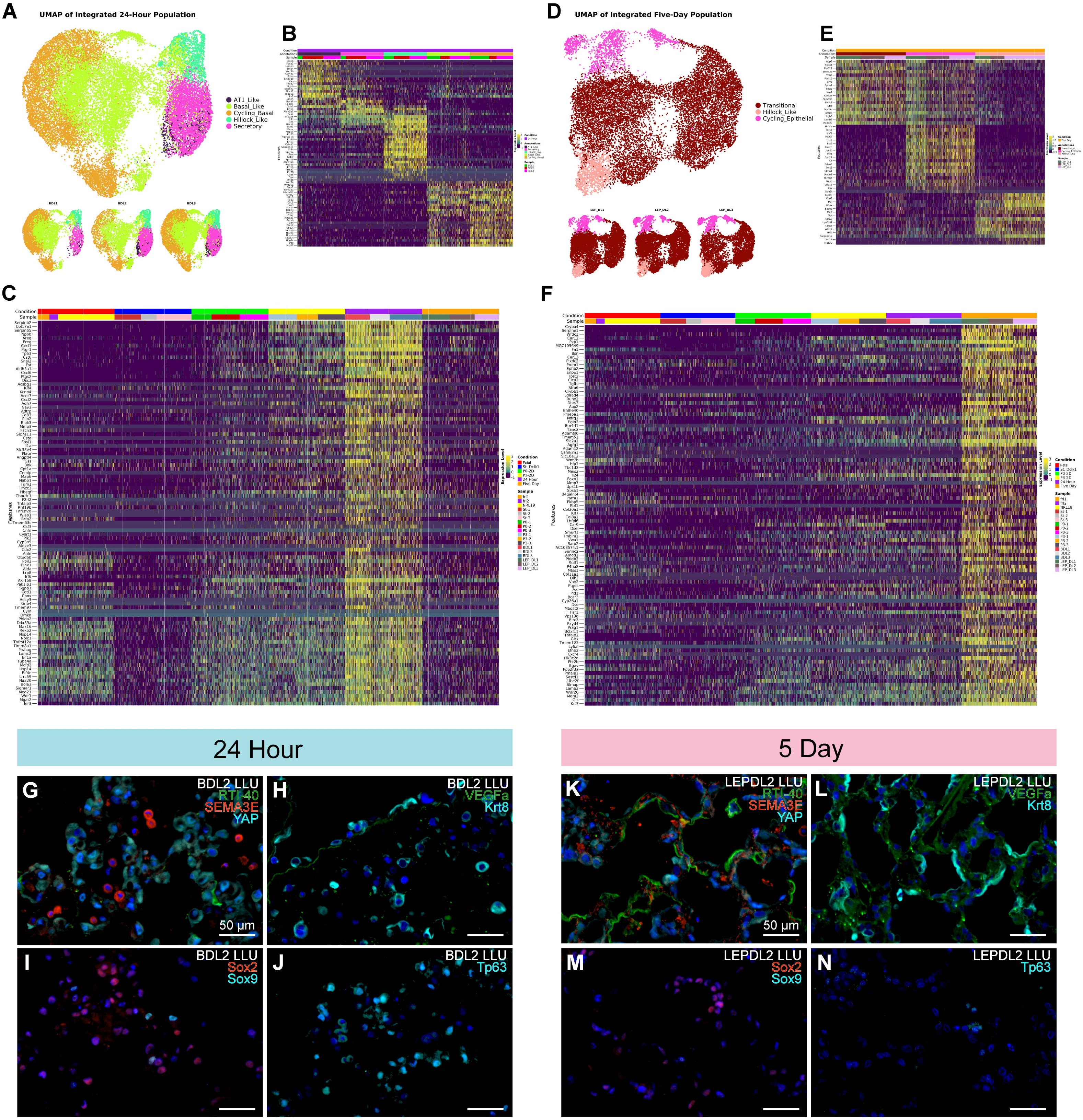


**Supplemental Figure 2.** 24-Hour and 5-day culture conditions profiled by scRNAseq and immunohistochemistry. A) Cross-sample UMAP embedding for the 24-Hour bioengineered lung condition showing annotated cell states and representation by each replicate. B) Heatmap showing genes marking each cell type cluster within the 24-Hour culture object. C) Heatmap showing genes marking the 24-Hour culture bioengineered lung condition against all other conditions in this study. D) Cross-sample UMAP embedding for the 5-Day bioengineered lung condition showing annotated cell states and representation by each replicate. E) Heatmap showing genes marking each cell type cluster within the 5-Day culture object. F) Heatmap showing genes marking the 5-Day culture bioengineered lung condition against all other conditions in this study. G-J) Representative immunofluorescence images from the left upper lobe (LLU) of the 24-hour bioengineered lungs (BDL2), showing expression of RTI-40, Sema3e, and YAP (G), Vegfa and Krt8 (H), proximal-distal lung development markers Sox9 and Sox2 (I) and basal cell marker, Tp63 (J). K-N) Matched immunostaining from the LLU of bioengineered lung (LEPDL2) at 5 days. Scale bars: 50 µm.

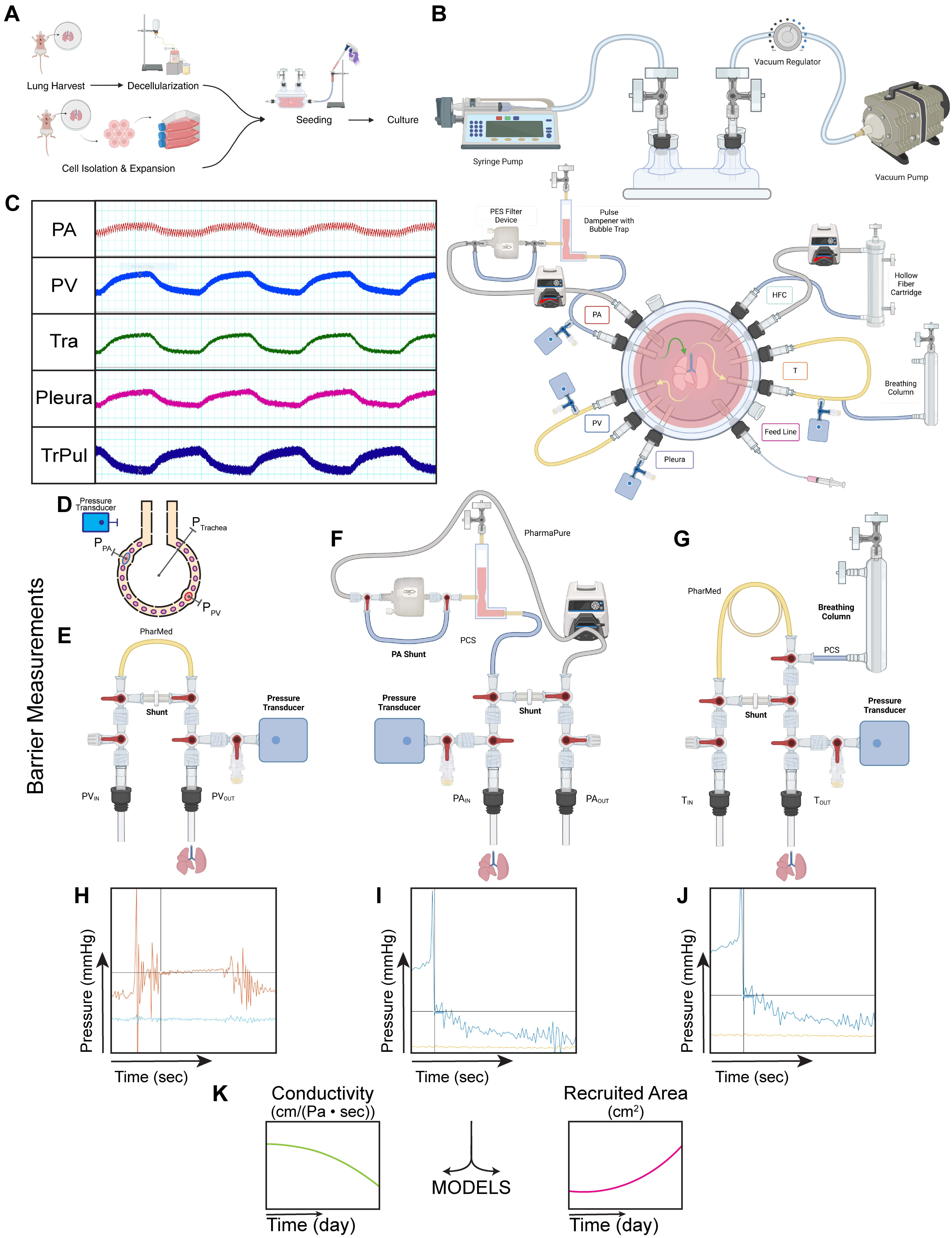


**Supplemental Figure 3.** Schematic of Bioengineering Methods. A) To create an engineered lung, a decellularized extracellular matrix scaffold is produced from an intact pair of explanted adult rat lungs, and cells are seeded into the airways for culture following the procedure described in Greaney & Raredon et al. 2024. B) The bioreactor apparatus consists of a sealed bioreactor with ports that allow for connections to external components to facilitate controlled and monitored media perfusion through the lungs at full cardiac output under biomimetic negative pressure with ventilation. C) Representative pressure-flow waveforms during culture. Pressure traces are recorded for the PA, PV, Trachea, and pseudo-Pleura, and a Transpulmonary pressure is calculated as the difference between the Pleura and Trachea pressures. D) Alveolar schematic showing fluid compartments. Media flows into the lung via the PA and out of the lung into the bioreactor chamber either through the PV , the trachea, or the visceral pleura. Pressures in end arterioles, end venules, and the alveoli of the lung was inferred mathematically using the method described in Engler et al 2019.
E) Pulmonary vein perfusion circuit. A small length of high-resistance tubing provides an optimized back pressure to stent the venous tree open as occurs in vivo.
F) Pulmonary artery perfusion circuit. Media is drawn from the bottom of the bioreactor basin by a peristaltic pump through a filter and hybrid pulse dampener and bubble trap device to prevent the vasculature from being compromised by debris or air bubble occlusions. G) Trachea perfusion circuit schematic. Media is allowed to flow out of the lung though the trachea due to a lack of barrier between the vasculature and airways, so this circuit allows control over the airway outflow with a length of resistance tubing and a breathing column allowing regulation of airway back-pressure and/or sterile opening to atmospheric pressure. H) PV Barrier Measurement. Example pressure trace and measurement selection for the barrier measurement of the PV at the initial stabilization of the instantaneous pressure reading. I) PA Barrier Measurement. Example pressure trace and measurement selection for the barrier measurement of the PA at the initial stabilization of the instantaneous pressure reading. J) Trachea Barrier Measurement. Example pressure trace and measurement selection for the barrier measurement of the PV at the initial stabilization of the instantaneous pressure reading. K) Schema showing modeling of fluidic conductivity per unit alveolar surface area and total recruited perfused alveolar in the lung, computed via the mathematical modeling approach described in Raredon et al. 2021. As barrier function increases, fluid conductivity through the capillary-alveolar barrier will decrease and the total recruited area of the lung, all else being equal, will increase.

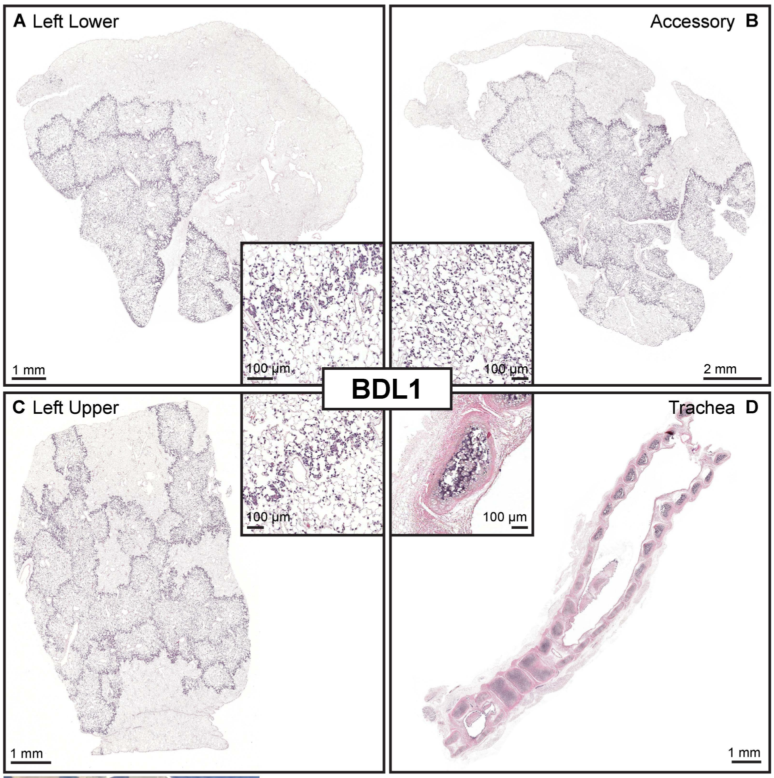


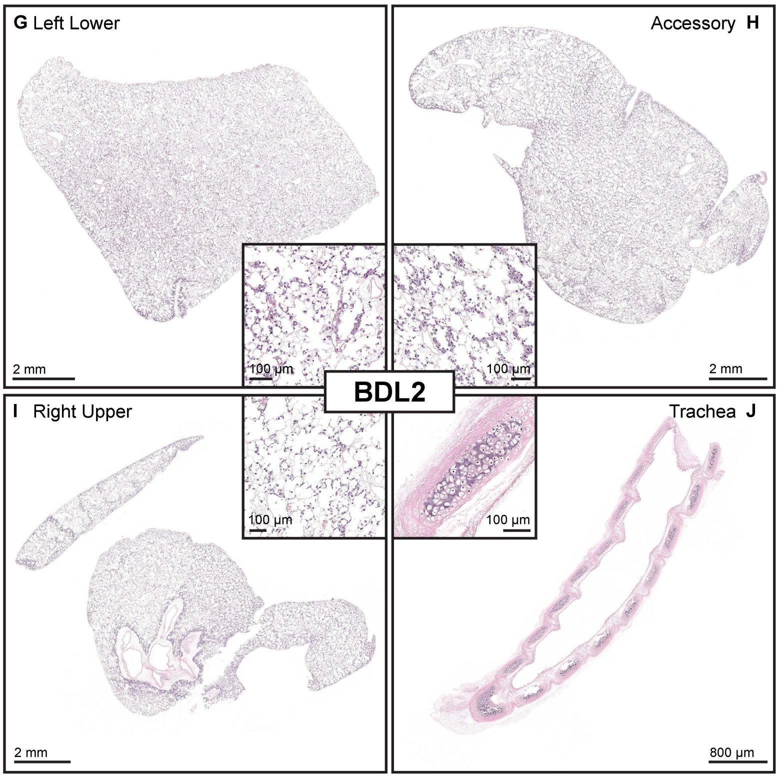


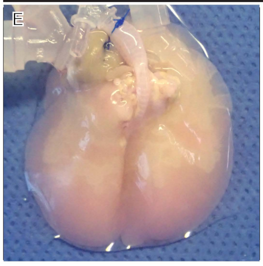


F

| Media Sampling Results | | | | | | |
|---|---|---|---|---|---|---|
| | Media | Day 1 | Day 2 | Day 3 | Day 4 | Day 5 |
| pH | 7.6 | 7.4 | | | | |
| Lactate | Low | 2.6 | | | | |
| Glucose | 313 | 289 | | | | |

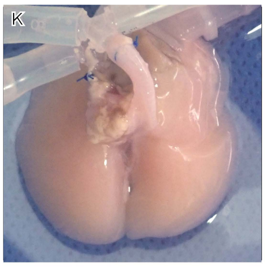


L

| Media Sampling Results | | | | | | |
|---|---|---|---|---|---|---|
| | Media | Day 1 | Day 2 | Day 3 | Day 4 | Day 5 |
| pH | 7.4 | 7.4 | | | | |
| Lactate | Low | 4.1 | | | | |
| Glucose | 302 | 280 | | | | |

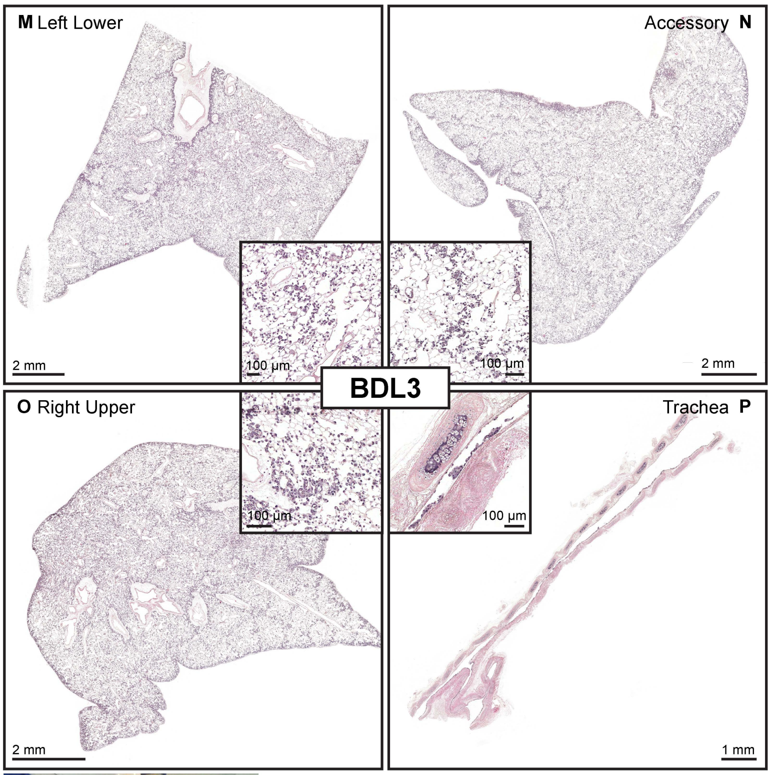


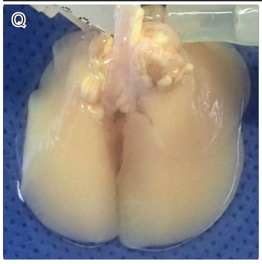


R

| Media Sampling Results | | | | | | |
|---|---|---|---|---|---|---|
| | Media | Day 1 | Day 2 | Day 3 | Day 4 | Day 5 |
| pH | 7.4 | 7.4 | | | | |
| Lactate | Low | 4.8 | | | | |
| Glucose | 302 | 268 | | | | |

**Supplemental Figure 4.** Histological and gross assessment of airway seeding across three independent bioengineered lung replicates after 24 hours of culture. A-D) BDL1 (Replicate 1) representative whole-mount sections stained with hematoxylin and eosin (H&E) from the left lower (A), accessory (B), left upper (C) and trachea (C), with higher magnification insets highlighting epithelial distribution. E) Gross image of BDL1 at the end of culture. F) Media sampling results for BDL1 at the start and end of culture, including measurements for pH, lactate and glucose. G-L) BDL2 (Replicate 2) representative H&E-stained sections from the left lower (G), accessory (H), right upper (I), and trachea (J), with corresponding magnified insets. K) Gross image of BDL2 at the end of culture. L) Media sampling results for BDL2 at the start and end of culture, including measurements for pH, lactate and glucose. M-P) BDL3 (Replicate 3) representative H&E-stained sections from left lower (M), accessory (N), right upper (O), and trachea (P) with insets. Q) Gross image of BDL3 at the end of culture. R) Media sampling results for BDL3 at the start and end of culture, including measurements for pH, lactate and glucose. Scale bars provided on all histological images.

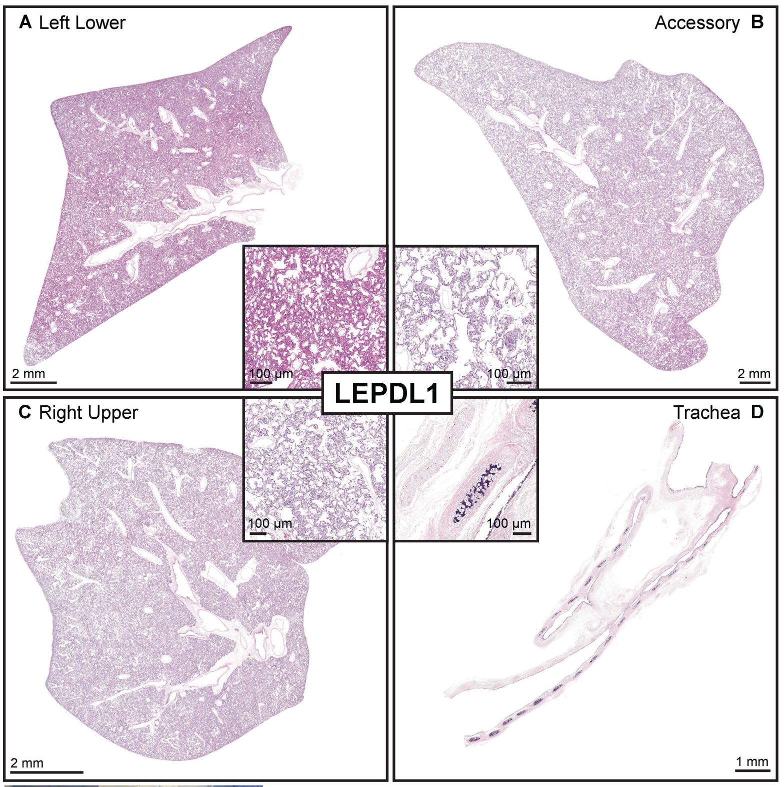


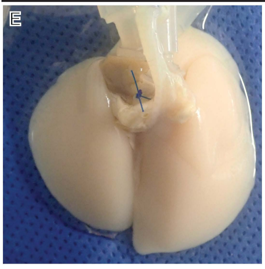


F

| Media Sampling Results | | | | | | |
|---|---|---|---|---|---|---|
| | Media | Day 1 | Day 2 | Day 3 | Day 4 | Day 5 |
| pH | 7.4/7.2 | 7.4 | 7.0 | 6.8 | 6.4 | 6.4 |
| Lactate | Low | 2.9 | 7.0 | 19.6 | 17.9 | 27.4 |
| Glucose | 288/264 | 282 | 236 | 101 | Low | 27 |

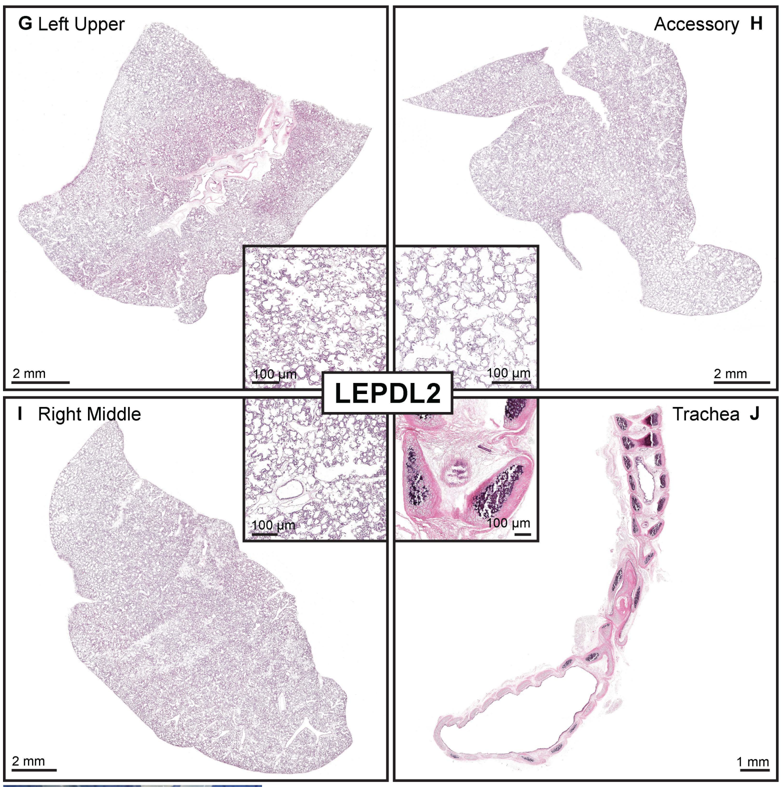


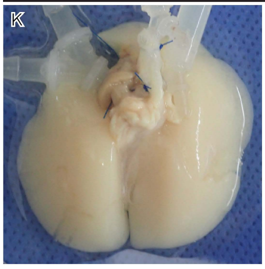


L

| Media Sampling Results | | | | | | |
|---|---|---|---|---|---|---|
| | Media | Day 1 | Day 2 | Day 3 | Day 4 | Day 5 |
| pH | 7.4/7.2 | 7.4 | 7.2 | 6.6 | 6.4 | 6.4 |
| Lactate | Low | 2.9 | 7.0 | 14.0 | 10.7 | 13.4 |
| Glucose | 314/272 | 262 | 257 | 154 | 88 | 93 |

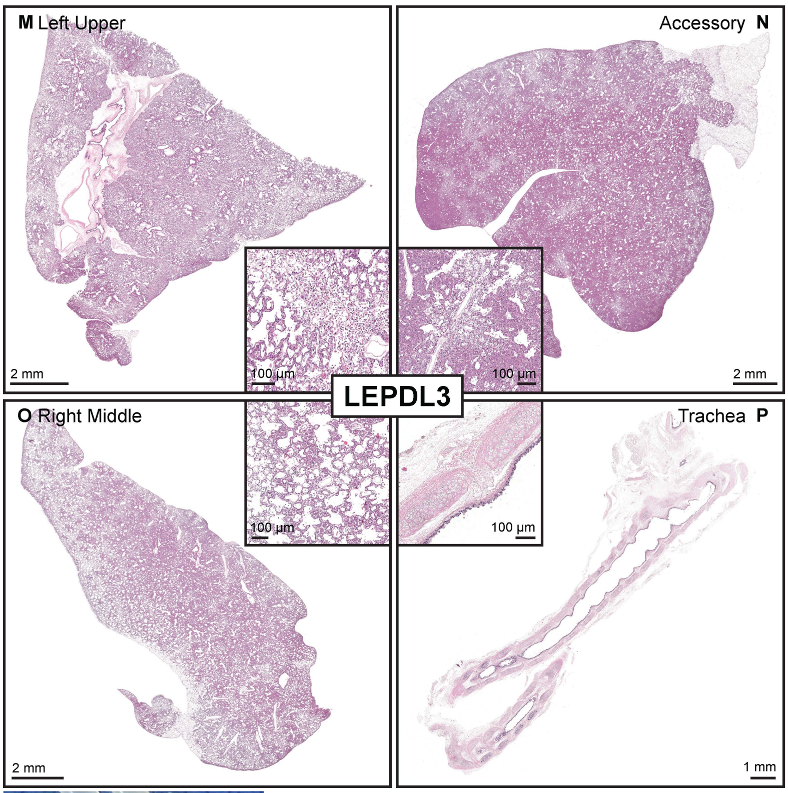


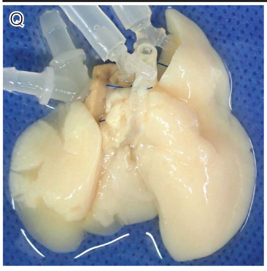


R

| Media Sampling Results | | | | | | |
|---|---|---|---|---|---|---|
| | Media | Day 1 | Day 2 | Day 3 | Day 4 | Day 5 |
| pH | 7.4/7.2 | 7.4 | 7.2 | 6.4 | 6.2 | 6.4 |
| Lactate | Low | 1.8 | 8.5 | 13.2 | 11.2 | 11.9 |
| Glucose | 282/272 | 278 | 242 | 98 | 57 | 63 |

**Supplemental Figure 5.** Histological and gross assessment of airway seeding across three independent bioengineered lung replicates after 5 days of culture. A-D) LEPDL1 (Replicate 1) representative whole-mount sections stained with hematoxylin and eosin (H&E) from the left lower (A), accessory (B), left upper (C) and trachea (C), with higher magnification insets highlighting epithelial organization within alveolar regions. LEPDL1 demonstrates broadly homogenous epithelial coverage, preserved tissue architecture, and flattening of epithelial cells within alveolar territories consistent with an ATI-like phenotype. E) Gross image of LEPDL1 at the end of culture. F) Media sampling results for LEPDL1 on each day of culture, including measurements for pH, lactate and glucose. G-J) LEPDL2 (Replicate 2) representative whole-mount sections stained with H&E from the left lower (G), accessory (H), left upper (I) and trachea (J), with higher magnification insets highlighting epithelial organization within alveolar regions. K) Gross image of LEPDL2 at the end of culture. L) Media sampling results for LEPDL2 on each day of culture, including measurements for pH, lactate and glucose. M-P) LEPDL3 (Replicate 3) representative whole-mount sections stained with H&E from the left upper (M), accessory (N), right middle (O) and trachea (P), with higher magnification insets. In contrast to LEPDL1-2, LEPDL3 displays spatially heterogenous epithelial coverage reflecting only partial recruitment during seeding. Poorly seeded and perfused regions show local cellular necrosis and tissue remodeling, whereas well-recruited regions display epithelial flattening and patent intrarectal capillary lumens resembling the morphology observed in LEPDL1 and LEPDL2. Q) Gross image of LEPDL3 at the end of culture. R) Media sampling results for LEPDL3 on each day of culture, including measurements for pH, lactate and glucose. Scale bars provided on all histological images.

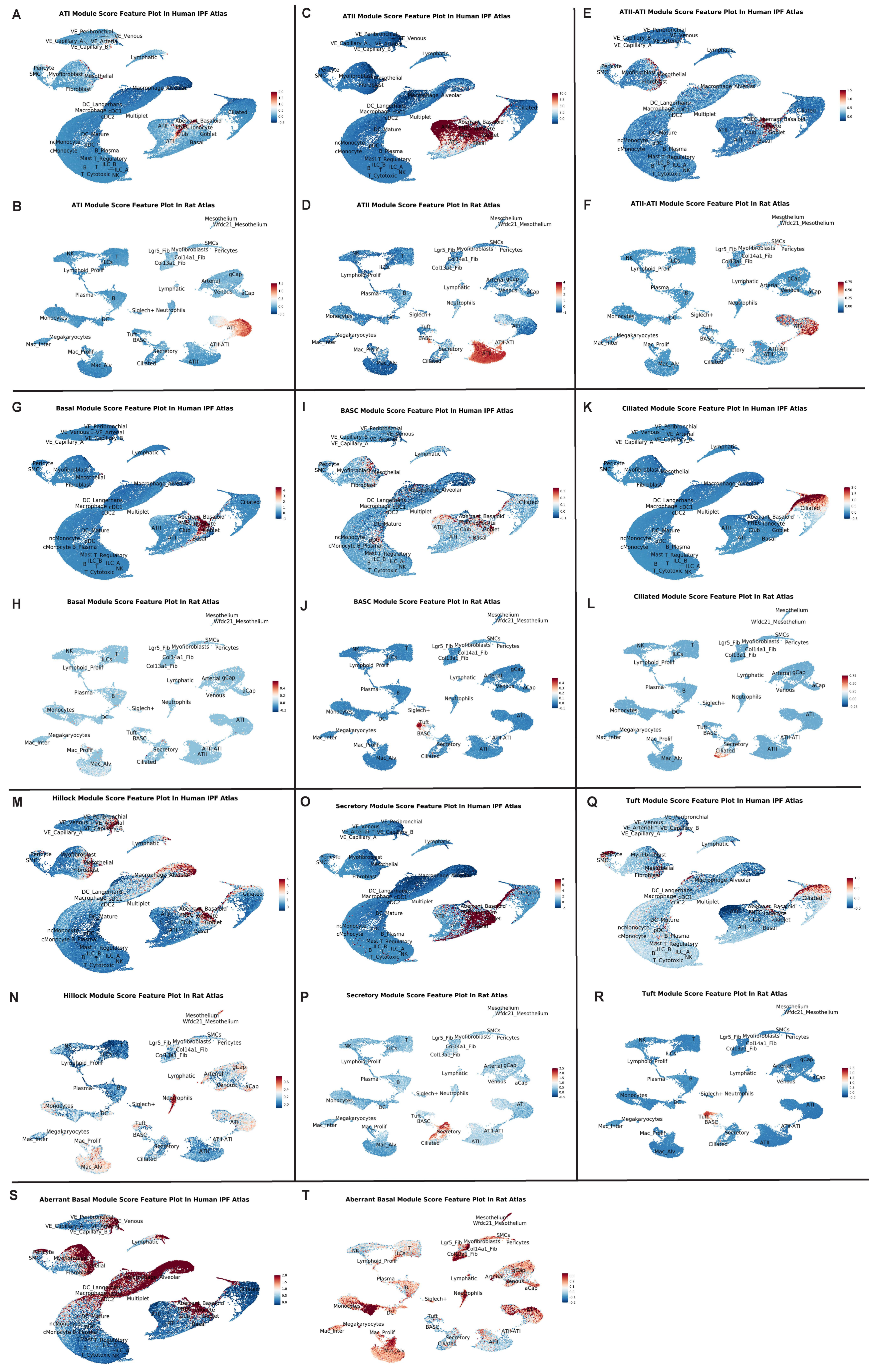


**Supplemental Figure 6.** Cross-species projection of epithelial module programs onto human IPF and nativerat lung atlases. A, B) ATI module scores in human IPF (A) and native rat (B) atlases. C, D) ATII module scores in human IPF (C) and native rat (D). E, F) ATII-ATI transitional module scores in human IPF (E) and native rat (F). G,H) Basal module scores in human IPF (G) and native rat (H). I, J) BASC module scores in human IPF (I) and native rat (J). K, L) Ciliated module scores in human IPF (K) and native rat (L). M, N) Hillock module scores in human IPF (Q) and native rat (R). S, T) Aberrant basal module scores in human IPF (S) and native rat (T).

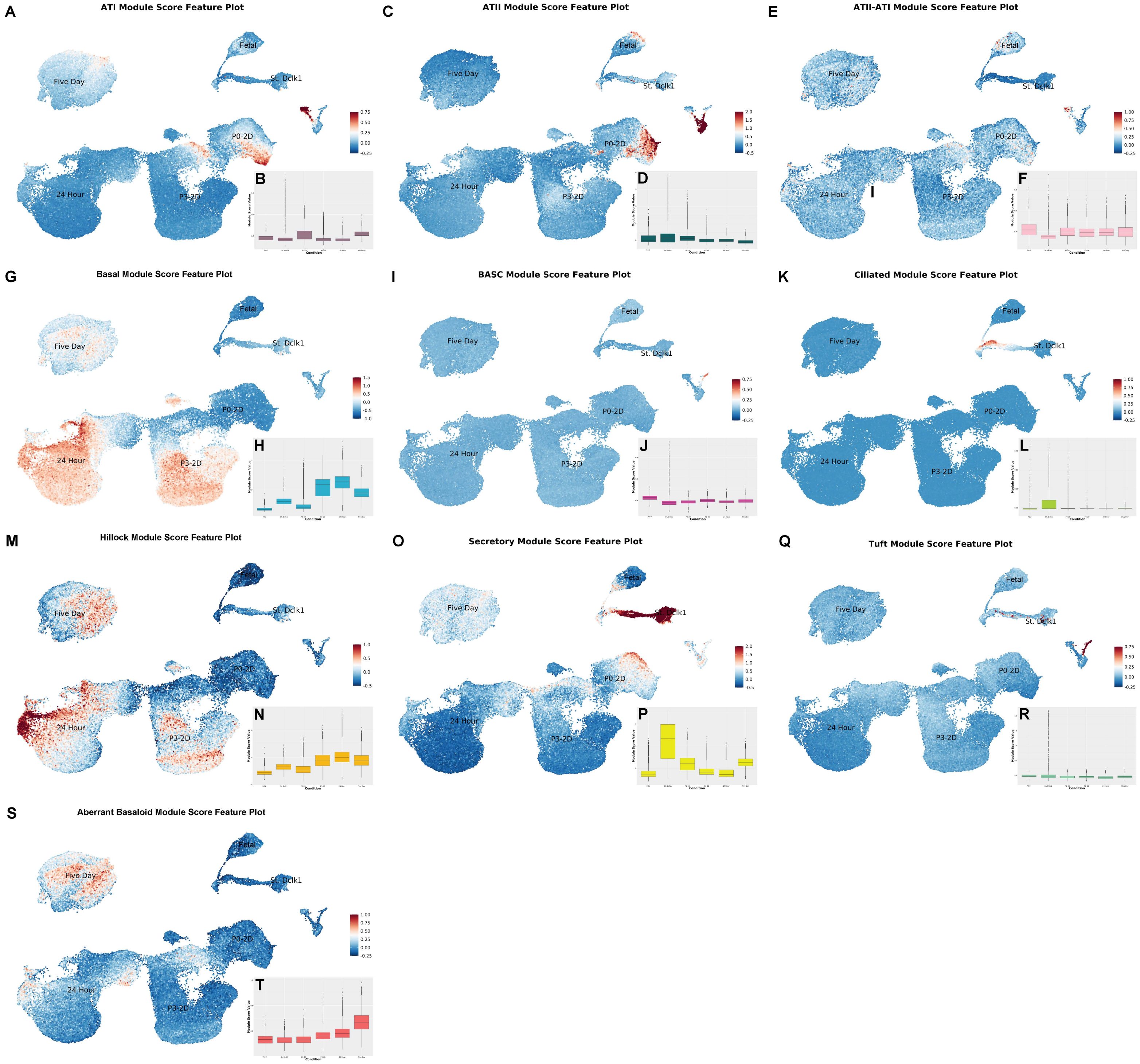


**Supplemental Figure 7.** Epithelial lineage and transitional state module scores across global object containing fetal, adult (St. Dclk1), P0-2D, P3-2D, 24 Hour and Five Day conditions. A, C, E, G, I, K, M, O, Q, S) UMAP feature plots showing per-cell module scores for ATI, ATII, ATII-ATI, basal, BASC, ciliated, secretory, tuft, hillock, and aberrant basaloid programs across Fetal, St. Dclk1, P0-2D, P3-2D, 24 Hour, and Five Day conditions. B, D, F, H, J, L, N, P, R, T) Corresponding box plots showing the distribution of module scores for the same programs across Fetal, St. Dclk1, P0-2D, P3-2D, 24 Hour, and Five Day conditions.

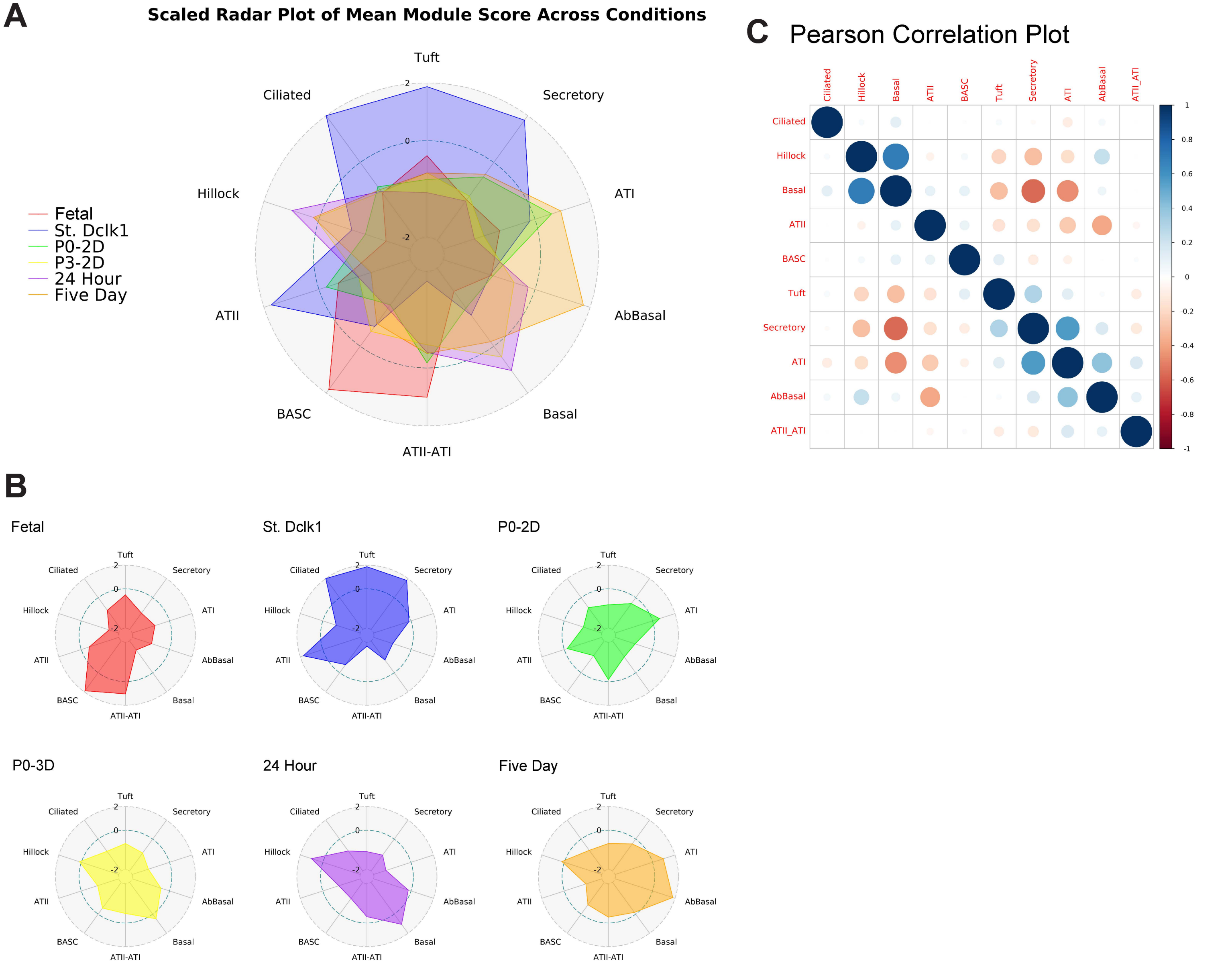


**Supplemental Figure 8.** Global organization and covariance of epithelial state modules across conditions. A) Scaled radar plot of mean module scores across all conditions with each axis representing an epithelial state program (ciliated, tuft, secretory, ATI, ATII, ATII-ATI, BASC, basal, aberrant basal, and hillock). Values are globally scaled to allow for direct comparison of relative module enrichment across conditions. B) Corresponding radar plots shown for each condition using the same global scaling as in (A) to visualize how epithelial state programs are redistributed across fetal, native, 2D, and 3D contexts. C) Pairwise correlation analysis computed across all cells profiled in this study, showing relationships between designated cellular state shifts. Note the high correlation between ATI-directed differentiation and the adoption of aberrant basaloid features, as well as the inverse relationship between ATI-directed differentiation and basal cell programming.

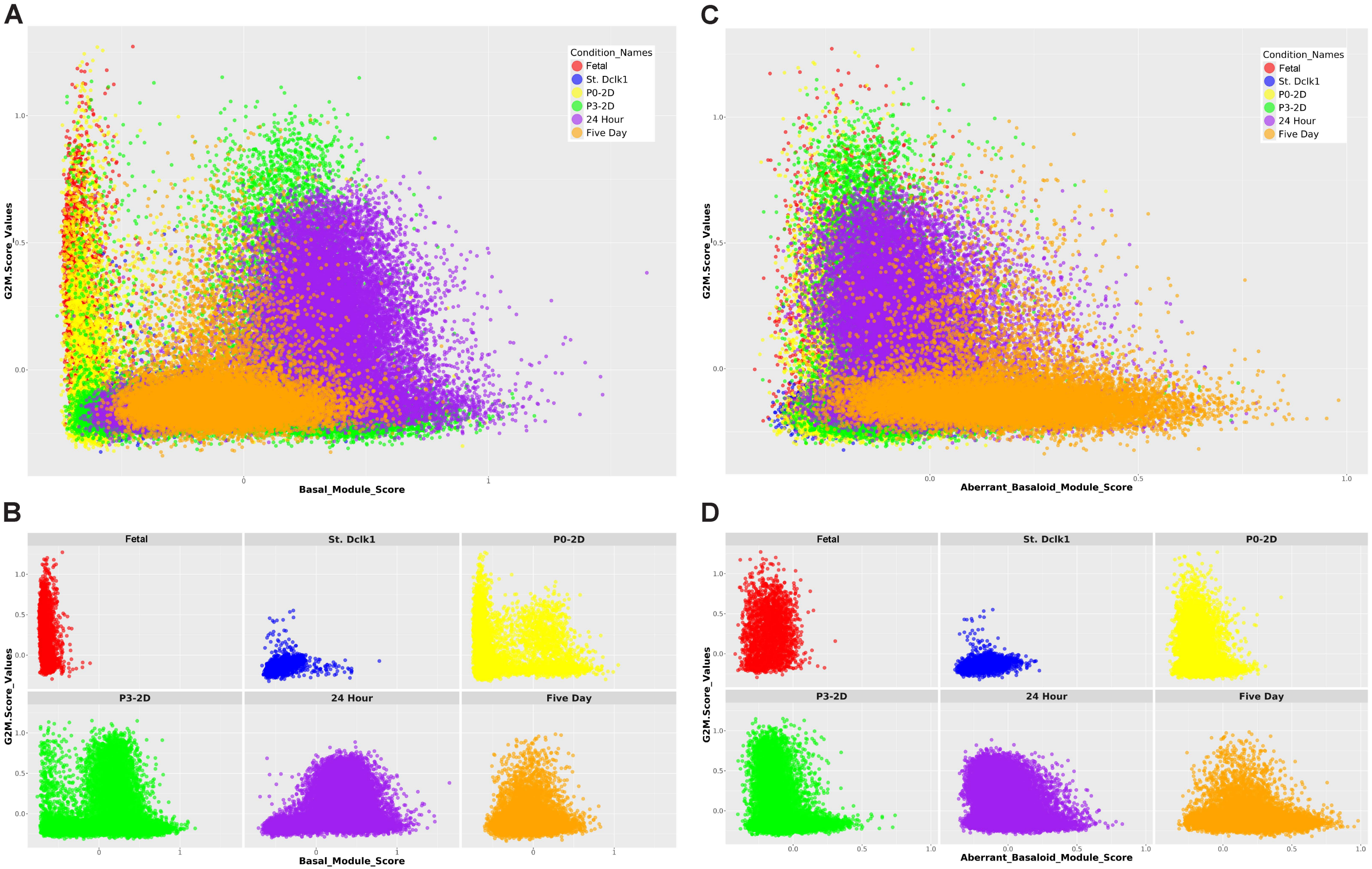


**Supplemental Figure 9.** Relationship between cell-cycle phase and basal-lineage module programs. A, B) Scatter plots showing G2M cell-cycle scores as a function of basal module scores for all cells (A) and stratified by experimental condition (B), indicating that canonical basal identity spans both cycling and non-cycling states. C, D) Corresponding plots showing G2M scores as a function of basaloid module scores for all cells (C) and by condition (D). Aberrant-basaloid cells appear to be preferentially distributed at low G2M values across conditions, consistent with reduced proliferative activity within this disease-associated epithelial program.
