## Supplemental Tables for "Epithelial Reprogramming and Transition during Pulmonary Bioengineering"

### Supplemental Table 1

| | Fetal-1 | Fetal-2 | Fetal-3 |
|---|---|---|---|
| Condition | Fetal | Fetal | Fetal |
| Sample Name | frl1 | frl2 | NRL19 |
| Library_Construction | 10x_v3 | 10x_v3 | 10x_v3 |
| Sequencing_Platform | Illumina NovaSeq 6000 | Illumina NovaSeq 6000 | Illumina NovaSeq 6000 |
| Sequencing_Date | 8/7/22 | 8/7/22 | 8/7/22 |
| Final_Cell_Number | 353 | 257 | 1705 |
| Mean_nFeature | 5620.286 | 5871.518 | 5056.74 |
| Mean_nCount | 41448.4 | 50035.67 | 34620.48 |
| Mean_percent.mt | 5.707461 | 5.426617 | 3.332762 |
| Reference_Genome | Rattus_norvegicus.Rnor_6.0.95 | Rattus_norvegicus.Rnor_6.0.95 | Rattus_norvegicus.Rnor_6.0.95 |

| | St-1 | St-2 | St-3 |
|---|---|---|---|
| Condition | Starting Dclk1 | Starting Dclk1 | Starting Dclk1 |
| Sample Name | iBASC_1 | iBASC_2 | iBASC_3 |
| Library_Construction | 10x_v3 | 10x_v3 | 10x_v3 |
| Sequencing_Platform | Illumina NovaSeq 6000 | Illumina NovaSeq 6000 | Illumina NovaSeq 6000 |
| Sequencing_Date | 3/8/24 | 6/29/24 | 7/12/24 |
| Final_Cell_Number | 1066 | 665 | 1434 |
| Mean_nFeature | 2560.987 | 2085.538 | 3810.317 |
| Mean_nCount | 14671.75 | 11063.82 | 28073.86 |
| Mean_percent.mt | 11.58867 | 10.80767 | 6.324766 |
| Reference_Genome | Rattus_norvegicus.Rnor_6.0.95 | Rattus_norvegicus.Rnor_6.0.95 | Rattus_norvegicus.Rnor_6.0.95 |

| | P0-1 | P0-2 | P0-3 |
|---|---|---|---|
| Condition | P0 | P0 | P0 |
| Sample Name | BASC11_P0 | BASC12_P0 | BASC7_P0 |
| Library_Construction | 10x_v3 | 10x_v3 | 10x_v3 |
| Sequencing_Platform | Illumina NovaSeq 6000 | Illumina NovaSeq 6000 | Illumina NovaSeq 6000 |
| Sequencing_Date | 9/13/24 | 9/13/24 | 5/7/24 |
| Final_Cell_Number | 2647 | 3937 | 4023 |
| Mean_nFeature | 3856.038 | 4731.501 | 4924.714 |
| Mean_nCount | 18023.8 | 35512.52 | 33240.29 |
| Mean_percent.mt | 9.916591 | 9.041633 | 8.213245 |
| Reference_Genome | Rattus_norvegicus.Rnor_6.0.95 | Rattus_norvegicus.Rnor_6.0.95 | Rattus_norvegicus.Rnor_6.0.95 |

| | P3-1 | P3-2 | P3-3 |
|---|---|---|---|
| Condition | P3 | P3 | P3 |
| Sample Name | BASC3_pre | BASC5_pre_SM | BASC6_pre_SM |
| Library_Construction | 10x_v3 | 10x_v3 | 10x_v3 |
| Sequencing_Platform | Illumina NovaSeq 6000 | Illumina NovaSeq 6000 | Illumina NovaSeq 6000 |
| Sequencing_Date | 5/7/24 | 3/20/24 | 3/21/24 |
| Final_Cell_Number | 8514 | 6356 | 8432 |
| Mean_nFeature | 3409.703 | 4599.608 | 3811.137 |
| Mean_nCount | 14316.37 | 28576.14 | 18348.35 |
| Mean_percent.mt | 6.305869 | 7.223156 | 6.662707 |
| Reference_Genome | Rattus_norvegicus.Rnor_6.0.95 | Rattus_norvegicus.Rnor_6.0.95 | Rattus_norvegicus.Rnor_6.0.95 |

| | BDL1 | BDL2 | BDL3 |
|---|---|---|---|
| Condition | Engineered 24 Hr | Engineered 24 Hr | Engineered 24 Hr |
| Sample Name | BDL1 | BDL2 | BDL3 |
| Library_Construction | 10x_v3 | 10x_v3 | 10x_v3 |
| Sequencing_Platform | Illumina NovaSeq 6000 | Illumina NovaSeq 6000 | Illumina NovaSeq 6000 |
| Sequencing_Date | 2/8/24 | 3/20/24 | 3/21/24 |
| Final_Cell_Number | 7430 | 6030 | 10128 |
| Mean_nFeature | 4441.572 | 4590.847 | 3293.004 |
| Mean_nCount | 24629.23 | 29158.79 | 14354.98 |
| Mean_percent.mt | 5.332731 | 9.263147 | 6.210431 |
| Reference_Genome | Rattus_norvegicus.Rnor_6.0.95 | Rattus_norvegicus.Rnor_6.0.95 | Rattus_norvegicus.Rnor_6.0.95 |

| | LEPDL1 | LEPDL2 | LEPDL3 |
|---|---|---|---|
| Condition | Engineered Five Day | Engineered Five Day | Engineered Five Day |
| Sample Name | LEP_DL1 | LEP_DL2 | LEP_DL3 |
| Library_Construction | 10x_v3 | 10x_v3 | 10x_v3 |
| Sequencing_Platform | Illumina NovaSeq 6000 | Illumina NovaSeq 6000 | Illumina NovaSeq 6000 |
| Sequencing_Date | 12/7/24 | 1/24/25 | 1/14/25 |
| Final_Cell_Number | 5571 | 5227 | 4930 |
| Mean_nFeature | 3299.115 | 2427.415 | 2168.454 |
| Mean_nCount | 19150.28 | 10013.01 | 8861.871 |
| Mean_percent.mt | 8.848041 | 8.666899 | 7.131733 |
| Reference_Genome | Rattus_norvegicus.Rnor_6.0.95 | Rattus_norvegicus.Rnor_6.0.95 | Rattus_norvegicus.Rnor_6.0.95 |

Supplemental Table 2 (page 1)

| Cell_Populations | Features |
|---|---|
| ATI_Genes | Pdpn, Aqp5, Akap5, Clic5, Cryab, Aebp1, Cyp2b1, Anxa8, Krt7, Cav1, Vegfa, Col4a4, Akap2, Rtkn2, Cav2, Sema3e, Icam1, Slc6a14, Timp3, Limch1, Cadm1, Cdkn2b, Flrt3, Ooep, Micu1, Cped1, Ehd2, Nckap5, Ccn2, Scnn1g, Arhgef26, Col4a5, Col12a1, Popdc2, Nnat, Ankrd1, Bdnf, Galnt18, Rab6b, Lamc2, Agrn, Col4a3, Fbln5, Tgfb2, Lox, Itga3, Cct6b, Lims2, Fzd2, Phyhipl, Wnt3a, Unc13d, Hbegf, Plekhh2, Schip1, Adgrv1, Sema6d, C1qtnf1, Zfp703, Trim3, Slc39a8, Wnt7a, Susd2, Sh3bp4, Stx11, Fstl3, Evala, RGD1310587, Shroom4, Slc16a12, Dusp26, Ptprr, Zfp365, Igfbpl1, Asic1, Plce1, Radil, Tmod1, Faim2, Rgs6, Zfp462, Ankrd29, Creb312, Rnf39 |
| ATII-ATI Genes | Edn3, Ajuba, Hspb2, Ydjc, Scx, Plau |
| ATII Genes | Defb3, Lamp3, S100g, Slc38a2, Defb4, Sftpb, Napsa, Fabp5 |
| Aberrant Basaloid Genes | Frmd5, Pld5, Gdf15, Cpa6, Pmepa1, Palld, Pxdn, Sox4, Ociad2, Cdh2, Ptgs2, Bicc1, Cst6, Kcns3, Tmem132d, Crlf1, Gas6, Cdkn2a, Myo3b, Pgbd5, Nkain4, Hmga2, Cank2n1, Pkhdl, Sema7a, Spink1, Nrk, Syt14, Baat, Pcp4, Prss1, Lhb, C5orf46, Capn6, Sox11, Crct1, Nog, Zbtb20, Trabd2b, Sdk1, Pthlh, Lpp, Ccbel, Bcat1, Ppplcb, Bpgm, Zpld1, Vat1, Isg15, Mdm2, Camk2n1, Mmp7, Ephb2, Vim, Tgfbi |
| Basal Genes | Krt5, Tp63, Wnt10a, Col17a1, Itga6, Comt, Dst |
| BASC Genes | Fbp2, Pou2f3, Sphkap, Tmem229a, Mia, Vil1, 1125, 1l17rb, Enpp6, Gfilb, Erich4, Bik, Cfc1, Asb15, Moxd1, Map2, Ddc, Chdh, Kcnj 13, St18, Pcdh20, Cwh43, Cdkn1c, Frrs1l, Slc22a15, Padi2, Trpm5, Mlc1, Crym, Col11a2, Mcub, Cdhr2, Sh2d7, Sct, Ampd3, Ank2, Elavl2, Zfp428, Arfgef3, Bmx, Rab3b, Anks4b, Eps8l3, Inava, Tmem51, RGD1562914, Arg2, Efs, Klhl23, Tubb2b, Ca11, Slc23a3, Tpd52\|1, Cldn6, Myo1a, Alox12, Abo, Mall, B3gnt8, Sel113, Egln3, Prox1, Spire2, Vsig2, Twist2, Nsg2, Etv4, Ascl2, Kif2c, Rhbdl2, Rab11fip4, Nrxn2, Entpd3, Cib2, Kcnk3, Ca6, Duoxa1 |
| Ciliated Genes | Ccdc40, Cfap161, Ccna1, AC103535.1, Fam92b, Ctrb1, RGD1560242, Armc4, Spag6l, Shc2, Tctex1d4, Pih1d3, Tsnaxip1, Daw1, RGD1564308, Spata18, RGD1308544, Drc7, Wdr38, Ttll, Ccdc24, Slc34a3, Tp73, AC128059.1, Dnaaf3, Stk33, Elmod1, Lrriq4, Ccdc103, Efcab1, Dnah5, Maats1, Stpg1, Vwa3b, Ccdc33, Mak, Xrral, Lkaaear1, RGD1560146, Ak8, Katnal2, Pex5l, Ccdc187, Lrrc73, Wdr93, MGC94891, Kcp, Ube2u, Lrrc9, Cfap157, Dpy1912, Dcdc2, Ccdc13, Rsph4a, Ubxn10 |
| Hillock Genes | Krt13, Prdm1, Grhl3, Scel, Emp1, Klf4, Cysrt1, MGC105649, Rhov, Paqr5, Dusp5, Pinlyp, Fxyd3, Gltp, Mxd1, Axna1, Arf6, S100a11, Junb, Tacstd2, S100a16, Prr13, Pdzk1ip1, Rab11a, Cd9, Axna7 |
| Secretory Genes | Scgb3a2, Bpifa5, Bpifb1, Scgb1a1, Tbz1, Ptpn13, Cpe, Gstm7, Cldn4, Pir |
| Tuft Genes | Rgs13, Gng13, Lrmp, Cd24, Rgs5, Ltc4s, Avil, Hmgn3, Hepacam2, Espn, Sox9, Nrgn, Tmem176a, Dclk1, AC120568.1, Akr783, Pstpip, Spib, Alox5, Atp2a3, Gnat3, Degs2, Bcl2l14, Camk2b, Pik3cg, Basp1, Siglec5, Rxrg, Itpr2, Hmx2 |

Supplemental Table 2 (page 2)

| | ATI | ATII_ATI | ATII | Aberrant_Basaloid | Basal | BASC | Ciliated | Hillock | Secretory | Tuft |
|---|---|---|---|---|---|---|---|---|---|---|
| ATI | SHARED | UNIQUE | UNIQUE | UNIQUE | UNIQUE | UNIQUE | UNIQUE | UNIQUE | UNIQUE | UNIQUE |
| ATII_ATI | UNIQUE | SHARED | UNIQUE | UNIQUE | UNIQUE | UNIQUE | UNIQUE | UNIQUE | UNIQUE | UNIQUE |
| ATII_ATI | UNIQUE | UNIQUE | SHARED | UNIQUE | UNIQUE | UNIQUE | UNIQUE | UNIQUE | UNIQUE | UNIQUE |
| Aberrant_Basaloid | UNIQUE | UNIQUE | UNIQUE | SHARED | UNIQUE | UNIQUE | UNIQUE | UNIQUE | UNIQUE | UNIQUE |
| Basal | UNIQUE | UNIQUE | UNIQUE | UNIQUE | SHARED | UNIQUE | UNIQUE | UNIQUE | UNIQUE | UNIQUE |
| BASC | UNIQUE | UNIQUE | UNIQUE | UNIQUE | UNIQUE | SHARED | UNIQUE | UNIQUE | UNIQUE | UNIQUE |
| Ciliated | UNIQUE | UNIQUE | UNIQUE | UNIQUE | UNIQUE | UNIQUE | SHARED | UNIQUE | UNIQUE | UNIQUE |
| Hillock | UNIQUE | UNIQUE | UNIQUE | UNIQUE | UNIQUE | UNIQUE | UNIQUE | SHARED | UNIQUE | UNIQUE |
| Secretory | UNIQUE | UNIQUE | UNIQUE | UNIQUE | UNIQUE | UNIQUE | UNIQUE | UNIQUE | SHARED | UNIQUE |
| Tuft | UNIQUE | UNIQUE | UNIQUE | UNIQUE | UNIQUE | UNIQUE | UNIQUE | UNIQUE | UNIQUE | SHARED |
